# An in-frame deletion mutation in the degron tail of auxin co-receptor *IAA2* confers resistance to the herbicide 2,4-D in *Sisymbrium orientale*

**DOI:** 10.1101/2021.03.04.433944

**Authors:** Marcelo R. A. de Figueiredo, Anita Küpper, Jenna M. Malone, Tijana Petrovic, Ana Beatriz T. B. de Figueiredo, Grace Campagnola, Olve B. Peersen, Kasavajhala V.S.K. Prasad, Eric L. Patterson, Anireddy S.N. Reddy, Martin F. Kubeš, Richard Napier, Christopher Preston, Todd A. Gaines

## Abstract

The natural auxin indole-3-acetic acid (IAA) is a key regulator of many aspects of plant growth and development. Synthetic auxin herbicides mimic the effects of IAA by inducing strong auxinic signaling responses in plants. Synthetic auxins are crucial herbicides in agriculture, made more important by the recent introduction of transgenic synthetic auxin resistant soybean and cotton. Currently, 41 weed species have evolved resistance to synthetic auxin herbicides and, in all but one case, the molecular basis of these resistance mechanisms is unknown. To determine the mechanism of 2,4-D resistance in a *Sisymbrium orientale* (Indian hedge mustard) weed population, we performed a transcriptome analysis of 2,4-D-resistant (R) and-susceptible (S) genotypes that revealed an in-frame 27-nucleotide deletion removing 9 amino acids in the degron tail (DT) of the auxin co-receptor Aux/IAA2 *(SoIAA2).* The deletion allele co-segregated with 2,4-D resistance in recombinant inbred lines. Further, this deletion was also detected in several 2,4-D resistant field populations of this species. Arabidopsis transgenic lines expressing the *SoIAA2* mutant allele were resistant to 2,4-D and dicamba. The IAA2-DT deletion reduced binding to TIR1 *in vitro* with both natural and synthetic auxins, causing reduced association and increased dissociation rates. This novel mechanism of synthetic auxin herbicide resistance assigns a new *in planta* function to the DT region of this Aux/IAA co-receptor for its role in synthetic auxin binding kinetics and reveals a potential biotechnological approach to produce synthetic auxin resistant crops using gene editing.

## INTRODUCTION

The natural auxin indole-3-acetic acid (IAA) regulates developmental processes in plant growth and morphogenesis. Synthetic auxin herbicides mimic the effects of IAA and induce strong changes in gene expression that ultimately lead to lethal plant growth responses. Auxin Response Factors (ARF) are transcription factors that bind to the promoter regions of auxin-responsive genes (Ulmasov et al., 1995). ARFs are regulated by transcriptional repressors called Aux/IAA proteins, which are rapidly degraded upon auxin binding by a class of SKP1/CULLIN1/F-BOX PROTEIN (SCF) protein complexes that contain an F-box protein coreceptor family called Transport Inhibitor Response 1/Auxin Signaling F-box (TIR1/AFB). Auxins act as a “molecular glue” to bring together SCF^TIR1/AFB^ and Aux/IAA. This process leads to Aux/IAA ubiquitination and subsequent degradation by the 26S proteasome, activating ARFs and the rapid transcription of auxin early responsive genes (Tan et al., 2007; Dinesh et al., 2016).

Aux/IAA proteins have four domains, including Domain I for transcriptional repression; Domain II consisting of the degron motif, a 13 amino acid sequence that binds to SCF^TIR1/AFB^ and auxin; and Domains III and IV called Phox and Bem1p (PB1 domain) respectively, which have homology to domains III and IV of ARFs forming homo and heterodimers that lead to transcriptional repression (Dinesh et al., 2015). Only the PB1 domain within the Aux/IAA structure has a defined crystal structure (Dinesh et al., 2015), while the other regions are characterized as intrinsically disordered regions (IDR). Mutations in certain regions of Aux/IAA genes including the degron can lead to auxin insensitivity, producing strong phenotypes with changes in leaf shape, plant size, root development, underdeveloped reproductive systems, and low seed production (Mockaitis & Estelle, 2008; Rinaldi et al., 2012). A mutation causing a Gly127Asn amino acid substitution in the degron motif of the *Aux/IAA16* gene in the weed species *Bassia scoparia* conferred robust resistance to the synthetic auxin herbicide dicamba (LeClere et al., 2018), but not cross-resistance to 2,4-dichlorophenoxyacetic acid (2,4-D) or fluroxypyr (Wu et al., 2021). In addition it caused substantial growth defects and reduced competitiveness (Wu et al., 2021), showing the consequential implications of mutations in the degron domains of Aux/IAA proteins. Currently, 41 different weed species have evolved resistance to synthetic auxin herbicides (Heap, 2020). Herbicide resistance mechanisms involve mutations in the gene encoding the target site protein(s) as well as non-target site changes to alter herbicide movement or rate of degradation to non-phytotoxic forms (Gaines et al., 2020). Synthetic auxin herbicide resistant weeds have been reported to have rapid metabolic degradation of the herbicide (Figueiredo et al., 2018), reduced herbicide translocation to the growing point (Goggin et al., 2016; Pettinga et al., 2018), and increased expression of transmembrane kinase and receptor proteins (Goggin et al., 2020). Except for the *IAA16* Gly127Asn degron mutation, the causative molecular basis of evolved synthetic auxin resistance in weeds is unknown (Gaines et al., 2020; Todd et al., 2020).

In 2005, a population of the weed species *Sisymbrium orientale* (Indian hedge mustard) was reported to be resistant to the synthetic auxin herbicides 2,4-D and 2-methyl-4-chlorophenoxyacetic acid (MCPA) in South Australia (Preston et al., 2013). In the resistant population, 2,4-D translocation was reduced (Dang et al., 2018) and progeny tests revealed that the resistance was inherited as a single dominant allele (Preston & Malone, 2015). Given the importance of synthetic auxins to agriculture, and the recent introduction of engineered resistance to 2,4-D and dicamba in soybean and cotton (Behrens et al., 2007; Wright et al., 2010), we investigated the mechanism of 2,4-D resistance in natural populations of the weed *S. orientale*. We performed a transcriptome analysis on Recombinant Inbred Lines (RILs) derived from a cross between 2,4-D-resistant (R) and -susceptible (S) genotypes and found an in-frame 27 bp deletion in the degron tail (DT) of the auxin co-receptor Aux/IAA2 *(SoIAA2).* This loss of nine amino acids due to DT deletion reduced binding of both natural and synthetic auxins to TIR1 and conferred cross-resistance to 2,4-D and dicamba, thus indicating the role of IAA2 in auxin perception and revealing the significance of the DT region in Aux/IAA degradation. These results demonstrate a novel resistance mechanism that confers field-evolved synthetic auxin herbicide resistance, advances our understanding of Aux/IAA domain functionality in auxin perception and signaling, and identifies a novel biotechnological tool that can be achieved by gene editing rather than transgene insertion for production of synthetic auxin resistant crops.

## RESULTS

### Identification of *IAA2* as a candidate gene for herbicide resistance through RNA-seq

The transcriptomes of untreated *S. orientale* plants were sequenced, consisting of six F4 RILs derived from a cross between 2,4-D resistant PB-R (Port Broughton-Resistant) and susceptible S (Cross A, 3 R-RILs and 3 S-RILs), and six F3 RILs from a cross between 2,4-D resistant PB-R2 and susceptible S (Cross B, 2 R-RILs and 4 S-RILs). Average reads per sample were 6.7×10^7^ with > 40 QC score. Only one transcript, *SoIAA2,* had differential expression between R-RILs and S-RILs of both PB-R and PB-R2 populations based on cutoff criteria of FDR <0.05, fold change of > |2|, and a consistent expression pattern in all R-RIL replicates compared to all S-RIL replicates. The IAA2 protein encoded by *SoIAA2* (3.3 fold lower expression in R than S, FDR<0.0001) showed a high sequence similarity to the Arabidopsis IAA2 protein encoded by AT3G23030 (Supplementary Fig. 1). No single nucleotide changes in any transcripts were identified that were shared among all R-RILs and different from all S-RILs. Inspection of the read alignments to *SoIAA2* identified a gap in read coverage for all R-RILs, suggesting a deletion in the R allele, whereas all S-RILs had continuous read coverage at this position (Supplementary Fig. 2). Transcriptome analysis led to the hypothesis that resistance was linked to this deletion mutation in *SoIAA2*.

### *SoIAA2* gene sequencing

A 239 bp region of the *SoIAA2* gene that includes the predicted deletion was amplified and sequenced from 3 R-RILs and 3 S-RILs. The 27 bp deletion in the R allele (gene and mRNA referred to as *SoIAA2_Δ27_*) was confirmed (Fig. 1A). The deletion results in an in-frame protein lacking nine amino acids (aa 73 to 81; protein referred to as *SoIAA2*_Δ9_) comprising most of the DT region between the degron and the PB1 domain (Fig. 1A). None of the S-RILs contained a deletion in *SoIAA2*, showing that the deletion was correlated with resistance.

**Figure 1.**
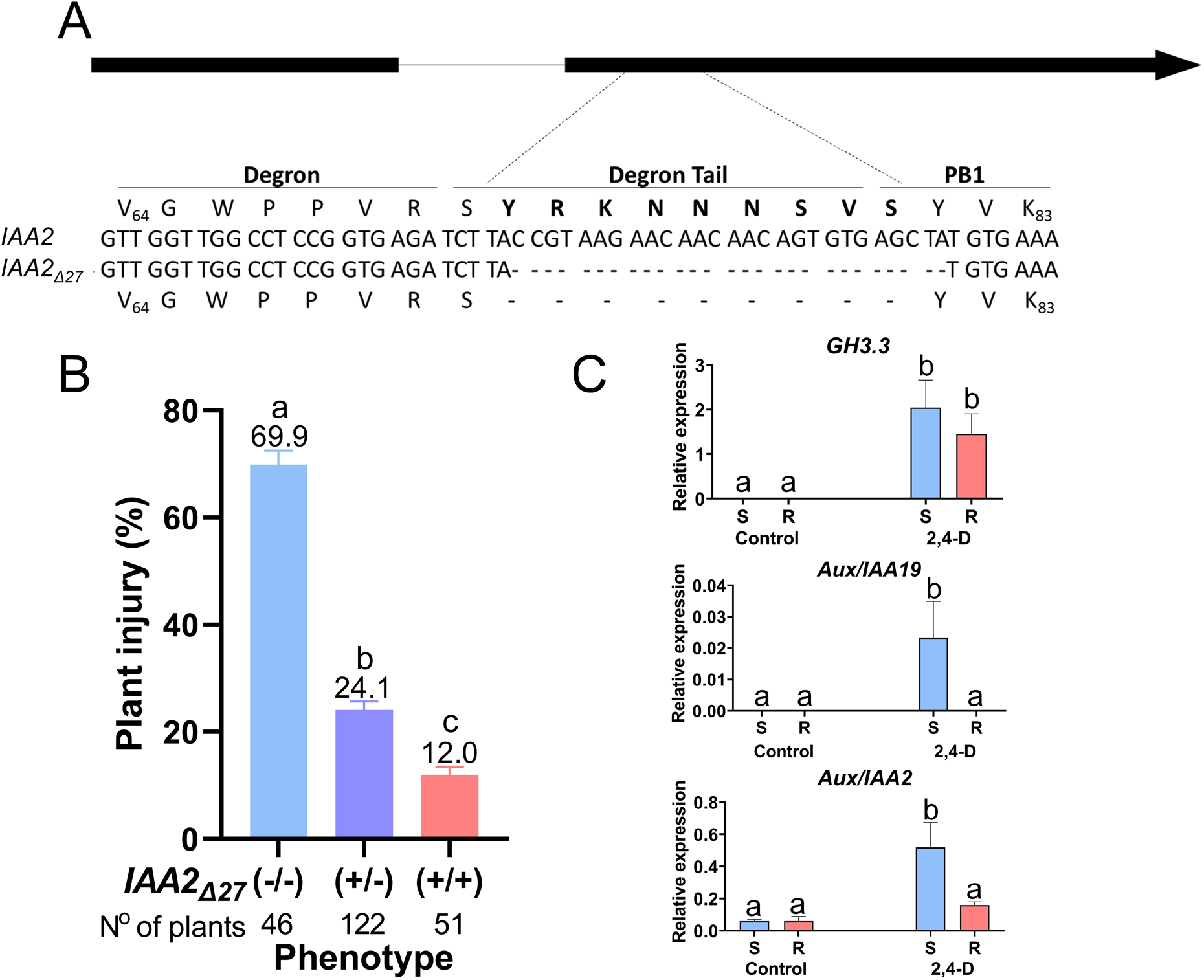
A deletion in *SoIAA2* confers resistance to 2,4-D. Deletion of 27 nucleotides from the *Sisymbrium orientale IAA2* gene results in a 9 amino acid deletion in the degron tail region of IAA2. A) *SoIAA2* schematic representation with two exons (black boxes) and one intron (line). Sequences showing differences between susceptible, S *(IAA2)* and resistant, R *(IAA2*_Δ27_) alleles, dashed lines show the nucleotides and amino acids that were deleted in the R allele. B) The deletion allele segregates with 2,4-D resistance in an F2 population. Plants were genotyped for the *IAA2* R and S alleles then sprayed with 250 gai ha^−1^ 2,4-D and evaluated for plant injury 28 d after treatment. Different letters indicate differences between means (p<0.05). C) Effect of 2,4-D treatment on genes related to auxin responses, *SoGH3.3; SoIAA19,* and *SoIAA2.* Herbicide treatment induces higher expression in S compared to R lines for *SoIAA19* and *SoIAA2* but not *SoGH3.3.* Different letters indicate differences between means (p<0.05). Plants were sprayed with 250 g ai ha^−1^ 2,4-D and gene expression was measured after 6 h using *cyclophilin* and *actin2* as normalization genes (n = 3 plants). Error bars correspond to standard deviation.

The presence of *SoIAA2*_Δ27_ in other 2,4-D resistant populations of *S. orientale* was investigated. Eight populations from South Australia, including parent populations PB-R and PB-R2 were screened for resistance to 2,4-D, with four populations identified as resistant. The *SoIAA2* gene was sequenced to determine the presence or absence of *SoIAA2*_Δ27_ in individual plants from each population (Supplementary Table 4). Full length *SoIAA2* was present in individuals from four susceptible populations P15, P31, P49 and P50. All individuals from the two parent resistant populations, PB-R and PB-R2, contained *SoIAA2*_Δ27_. Individuals from resistant population P17 were found to be homozygous for *SoIAA2*_Δ27_, or heterozygous for *SoIAA2*_Δ27_ and WT *SoIAA2*, suggesting a segregating population for this locus. Population P28, although resistant, did not contain *SoIAA2*_Δ27_, suggesting a different synthetic auxin herbicide resistance mechanism may have evolved in this population. Overall, sequence data identified a 27-base pair deletion in *SoIAA2* as the candidate mutation associated with resistance to 2,4-D in several *S. orientale* populations from Port Broughton, Australia.

### Segregation of *SoIAA2*_Δ27_

The heritable association between *SoIAA2*_Δ27_ and the resistance phenotype was confirmed by forward genetics through segregation and genotyping analysis. Susceptible progeny of a self-pollinated heterozygous individual (70% visual injury after 250 g ai ha^−1^ 2,4-D treatment) were homozygous for WT *SoIAA2;* heterozygous offspring showed an average of 24% injury; and resistant plants (average 12% injury) were homozygous for *SoIAA2*_Δ27_ (Fig. 1B). Therefore, the resistance phenotype segregates with the mutant allele *SoIAA2*_Δ27_. Relative expression of auxin responsive genes *IAA2, IAA19,* and *GH3.3* was compared between R and S plants after 250 g ai ha^−1^ 2,4-D treatment. In untreated plants, *GH3.3* and *IAA19* had no detectable transcripts for both genotypes while *SoIAA2* had low expression (Fig. 1C). Both S and R plants had similar elevation of *GH3.3* expression levels after 2,4-D treatment; however, expression levels of both *IAA19* and *IAA2* showed an increase in S compared to R (Fig. 1C).

### Expression of *SoIAA2*_Δ24_ in Arabidopsis confers 2,4-D and dicamba resistance

To test if *SoIAA2*_Δ27_ was sufficient to confer 2,4-D resistance, Arabidopsis was transformed with an empty *pFGC5941* vector (∅), *SoIAA2,* and *SoIAA2*_Δ27_ under the *CaMV35S* promoter (Supplementary Fig. 3). Genotypes were confirmed by PCR and plants containing the empty vector and WT *SoIAA2* did not show differences in plant phenotype (Fig. 2A). Arabidopsis transformed lines that were heterozygous for *SoIAA2*_Δ27_ had lanceolate shape leaves and a minor reduction in plant size. Plants homozygous for *SoIAA2*_Δ27_ had strong morphological abnormalities, dwarfism, a reduced number of reproductive organs, and lower seed production (Fig. 2A).

**Figure 2.**
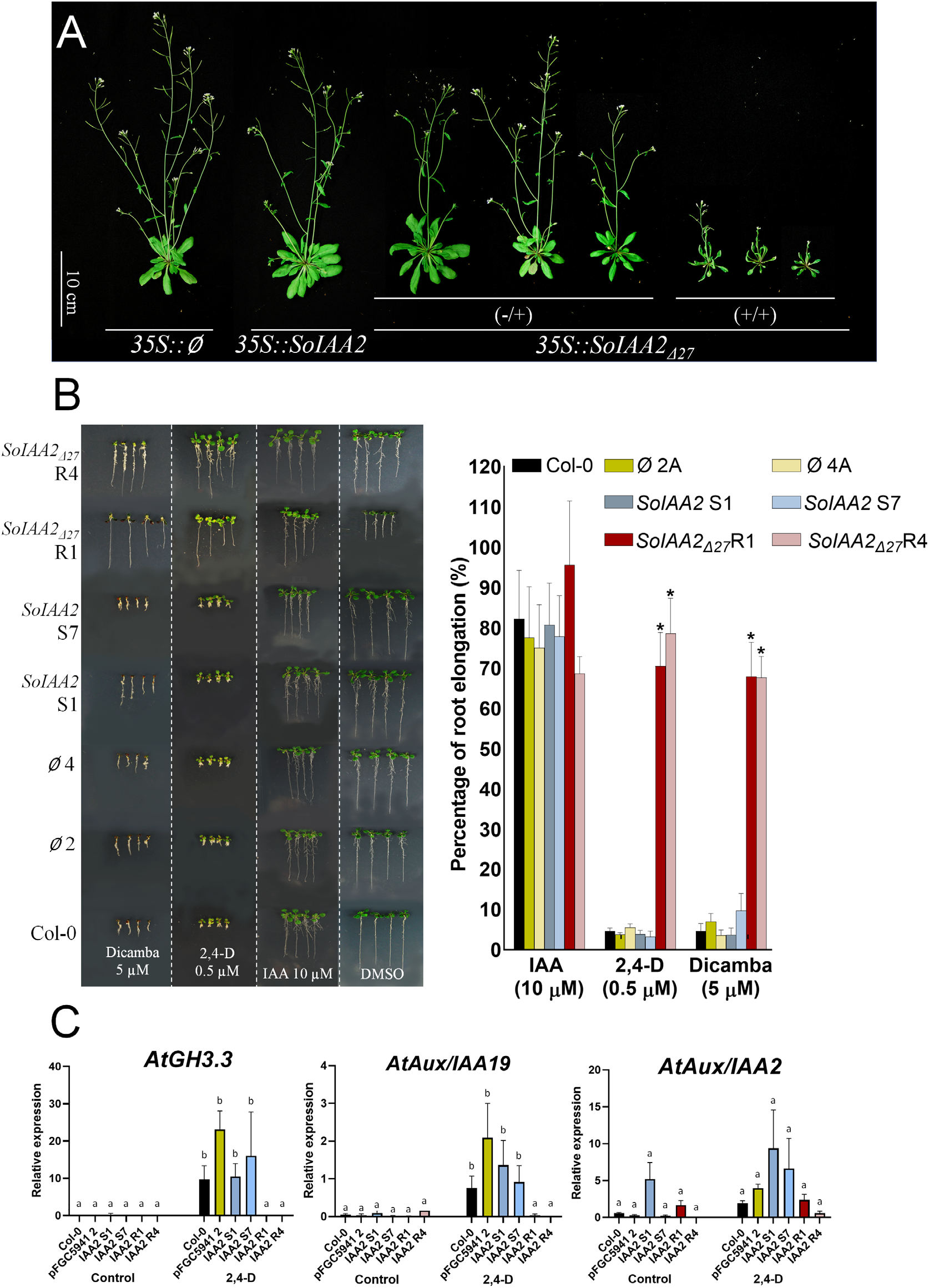
Transformation of *Arabidopsis thaliana* with the *IAA2* wild-type allele and *IAA2*_Δ27_ allele from *Sisymbrium orientale.* Treatments with the auxin herbicides 2,4-D and dicamba result in different root growth and gene expression responses consistent with the hypothesis that the *IAA2*_Δ27_ allele confers resistance. A) Pictures of transgenic lines 28 d after germination. Lines with vector control (35S:: ∅), *IAA2* (35S:: *So*IAA2) and *IAA2*_Δ27_ (35S::*So IAA*_Δ27_). The symbols +/− correspond to heterozygous and +/+ correspond to homozygous plants. Pictures are representative of at least three transformed lines selected for glufosinate resistance. B) Only the *IAA2*_Δ27_ allele grows on media containing auxin herbicides. Left panel, representative photos of seedlings of Col-0 (WT) and two independent lines for each (∅), *IAA2* and *IAA*_Δ27_ constructs growing on agar plates of auxins or the same final concentration of DMSO as control, 16 d after germination. Right panel, root elongation data. Error bars correspond to standard deviation. Asterisks correspond to statistical significances between different treatments (n ≤ 12 plants; p-value ≤ 0.05). C) Gene expression of auxin responsive genes *AtGH3.3* and *AtAux/IAA19* increased at 6 h after 2,4-D post-emergence treatment in Arabidopsis Col-0, empty vector (pFGC5941 2), and transgenic lines expressing wild-type *IAA2 (35S::SoIAA2; SoIAA2* S1 and S7), while no gene expression increase occurred in transgenic lines expressing *IAA2*_Δ27_ *(35S::SoIAA*_Δ27_; *SoIAA2* R1 and R4) (n=3). Different letters indicate differences between means (for *AtGH3.3,* p<0.05; for *AtAux/IAA19,* p<0.1) and error bars correspond to standard deviation. No significant gene expression changes occurred for *AtAux/IAA2* 6 h after 2,4-D post emergence treatment.

A root growth assay in the presence and absence of IAA and synthetic auxins was performed using the transformed Arabidopsis lines. Col-0 (WT) and plants containing the null vector (∅2 and ∅4) and *SoIAA2* (S1 and S7) were susceptible to inhibition by 2,4-D (0.5 μM) and dicamba (5 μM), with 90% inhibition of root elongation compared to the DMSO control (Fig. 2B). Arabidopsis transformed with *SoIAA2_Δ27_* (lines R1 and R4) exhibited high resistance to 2,4-D and dicamba, showing inhibition of root growth of only 20 to 30% compared to the DMSO control (Fig. 2B). No differences in root elongation were observed between any of the Col-0 or transgenic lines with the natural auxin IAA at 10 μM. *SoIAA2_Δ27_* expressing lines were also resistant to 2,4-D and dicamba at the whole plant level with foliar herbicide applications (Supplementary Fig. 4). The auxin responsive genes *AtIAA19* and *AtGH3.3* exhibited an increase in expression levels following 2,4-D foliar treatment in Col-0, null vector (Ø2), and *SoIAA2* (S1 and S7) lines, while lines transformed with *SoIAA2_Δ27_* (R1 and R4) showed no increase in expression of *AtIAA19* and *AtGH3.3* following 2,4-D treatment (Fig. 2C). *AtIAA2* expression was similar to that of Col-0 and null vector lines. Taking all these results together, *SoIAA2_Δ27_* is sufficient to confer resistance to both 2,4-D and dicamba when expressed in Arabidopsis.

### *So*IAA2 and *So*IAA2_Δ9_ protein structure

The Aux/IAA DT has previously been shown to be critical for the Aux/IAA interaction with TIR1, facilitating the ubiquitination of the PB1 domain (Niemeyer et al., 2020). Compared to Aux/IAA proteins, *So*IAA2 has a shorter and relatively structured DT. The nine amino acid deleted section of the DT is hydrophilic and *in silico* models predict the deleted DT region and the PB1 domain to be predominantly structured (Supplementary Fig. 5). To examine structural changes induced by the deletion mutation, full length *So*IAA2 and *So*IAA2_Δ9_ were expressed and purified to perform circular dichroism (CD) spectrometry and size exclusion chromatography.

Size exclusion chromatography showed that both mutant and wild-type proteins purify as oligomers of varied composition depending on protein concentration (Supplementary Fig. 6), which has also been seen for other Aux/IAA proteins (Kim et al., 1997; Nanao et al., 2014). While the deletion did not disrupt or enhance the oligomer formation, it did enhance the overall stability of the protein. During the purification process, the wild type protein displayed a much lower concentration threshold than the deletion mutation and could not be purified in a state higher than a predicted hexamer, whereas the deletion mutation was easily purified above an estimated decamer. CD spectrometry data are nearly identical for both proteins regardless of oligomeric state, although the concentration range needed for accurate CD data collections places the protein in a tetramer-hexamer state based on the gel filtration column elution profile data. A BeStSel analysis of the data (Micsonai et al., 2018) results in an alpha:beta:other content of 13%: 38%: 49%, which agrees with a partial structure from the PB1 domain and about half of the protein being disordered or “other”. This suggests the 9 amino acid deletion does not impact the overall folding of the protein (Supplementary Fig. 6).

### *So*IAA2 and *So*IAA2_Δ9_ affinity binding analysis

To determine the mechanism by which the DT deletion in *So*IAA2 interferes with synthetic auxin herbicide binding, a surface plasmon resonance (SPR) assay was performed (Lee et al., 2014). The assay measures the auxin-induced assembly of TIR1/AFB receptors with degron peptides, which is representative of the co-receptor complex (Calderón Villalobos et al., 2012). We used a biotinylated 24 aa peptide (*So*IAA2) consisting of the degron core (9 aa), DT (9 aa), and a small fraction of PB1 (6 aa); and a biotinylated 15 aa peptide lacking the DT segment (*So*IAA2_Δ9_) (Supplementary Table 3). Both peptides bound to TIR1 and to AFB5 in the presence of IAA (Fig. 3A and 3B, respectively). However, the assay showed weaker binding by *So*IAA2_Δ9_ to TIR1 with IAA compared to WT *So*IAA2, and similarly weaker binding with both synthetic auxins 2,4-D and dicamba (Fig. 3A; Supplementary Table 5). Binding kinetic experiments showed that weaker binding translates as lower affinity (a higher equilibrium dissociation constant, K_D_) for *So*IAA2_Δ9_ than for *So*IAA2 in each TIR1-auxin complex (K_D_ IAA=41 vs 11 nM; 2,4-D 300 vs 140 nM; dicamba 700 vs 250 nM) (Fig. 3; Supplementary Table 5). The association rate constants (*k*_a_) of *So*IAA2_Δ9_ were always lower than for WT *So*IAA2 in association with TIR1 (Supplementary Table 5). The dissociation rate constants (*k*_d_) were higher for the TIR1-*So*IAA2_Δ9_ complex with IAA and dicamba, but the *k*_d_ was slightly lower for 2,4-D with the TIR1-SoIAA2_Δ9_ complex (Supplementary Table 5). The kinetic data suggest that the lower affinity binding of the mutant will be slower to form and be less stable in the co-receptor complex in the presence of both auxin and synthetic auxin herbicides, resulting in reduced ubiquitination, longer lifetimes of Aux/IAA proteins, and reduced sensitivity to auxins.

**Figure 3.**
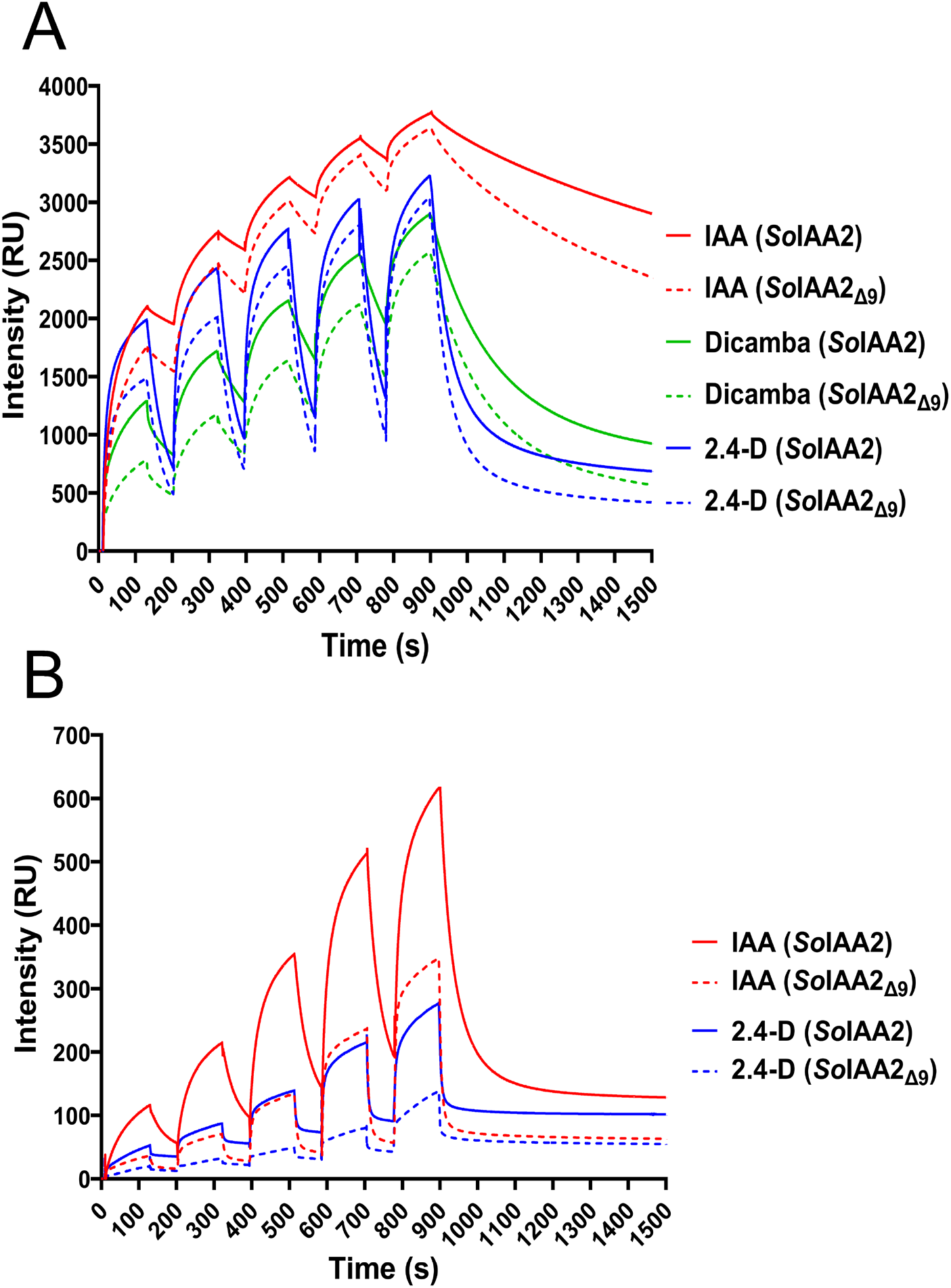
The mutant IAA2_Δ9_ binds to the receptors TIR1 and AFB5 with poorer affinity than the WT IAA2. Surface plasmon resonance was used to measure the binding of purified receptor protein to each degron peptide using single cycle kinetics. (A) Typical sensorgrams showing binding and dissociation of TIR1 with each degron peptide at a series of rising concentrations of each auxin. The same analysis with AFB5 (B), which characteristically displays faster kinetics for both association of the complex, and dissociation. Auxins were mixed with the TIR1 or AFB5 preparation in advance of injection over the biotinylated IAA2 peptides using standard Biacore double baseline subtraction with single cycle kinetic routines. RU, resonance units.

### Protein model in Pymol

A docking study was performed using the structures of the PB1 domain of *Ps*IAA4 and the ASK-TIR1-2,4-D-degron complex to explore the role of the DT for the association between TIR1 and IAA2. The PB1 and the DT were manually adjusted according to the TIR1 interaction of Niemeyer et al. (2020). The nine residue deletion shortened the DT by approximately 28 Å (Fig. 4), which in turn reduced the surface area of the interaction between the IAA2 and TIR1 proteins, loosening interactions with leucine-rich repeats 3-6 on the TIR1 protein even though the core degron remained in its binding site. This foreshortening would likely prevent the simultaneous binding of PB1 at its binding site located at the side of TIR1 and the core degron in its binding pocket on the top of TIR1 (Fig. 4, 5). Taking this observation in combination with the reduced binding affinity recorded by SPR, we propose a model to explain why the deletion in the *SoIAA2* DT results in resistance to auxin herbicides. Based on the protein-protein interaction model created by Niemeyer et al. (2020), in the presence of auxin the WT version of *So*IAA2 binds to TIR1 by interacting with the degron and the PB1 domain, possibly with the KR motif associating at the opposite pole of TIR1 to PB1 (Fig. 5). In this conformation, the IDR is displayed across the top surface of TIR1 with all lysine residues accessible for ubiquitination. In *So*IAA2_Δ9_ resistant individuals, the reduction of the molecular distance between the core degron and the PB1 domain precludes simultaneous binding at both sites. In combination with lower overall affinity for *So*IAA2_Δ9_, the lifetime of the complex is reduced and, hence, the chance of polyubiquitination is reduced. The loss of a lysine within the DT also reduces the number of available ubiquitination sites and this may also contribute to stabilizing the mutant *So*IAA2_Δ9_.

**Figure 4.**
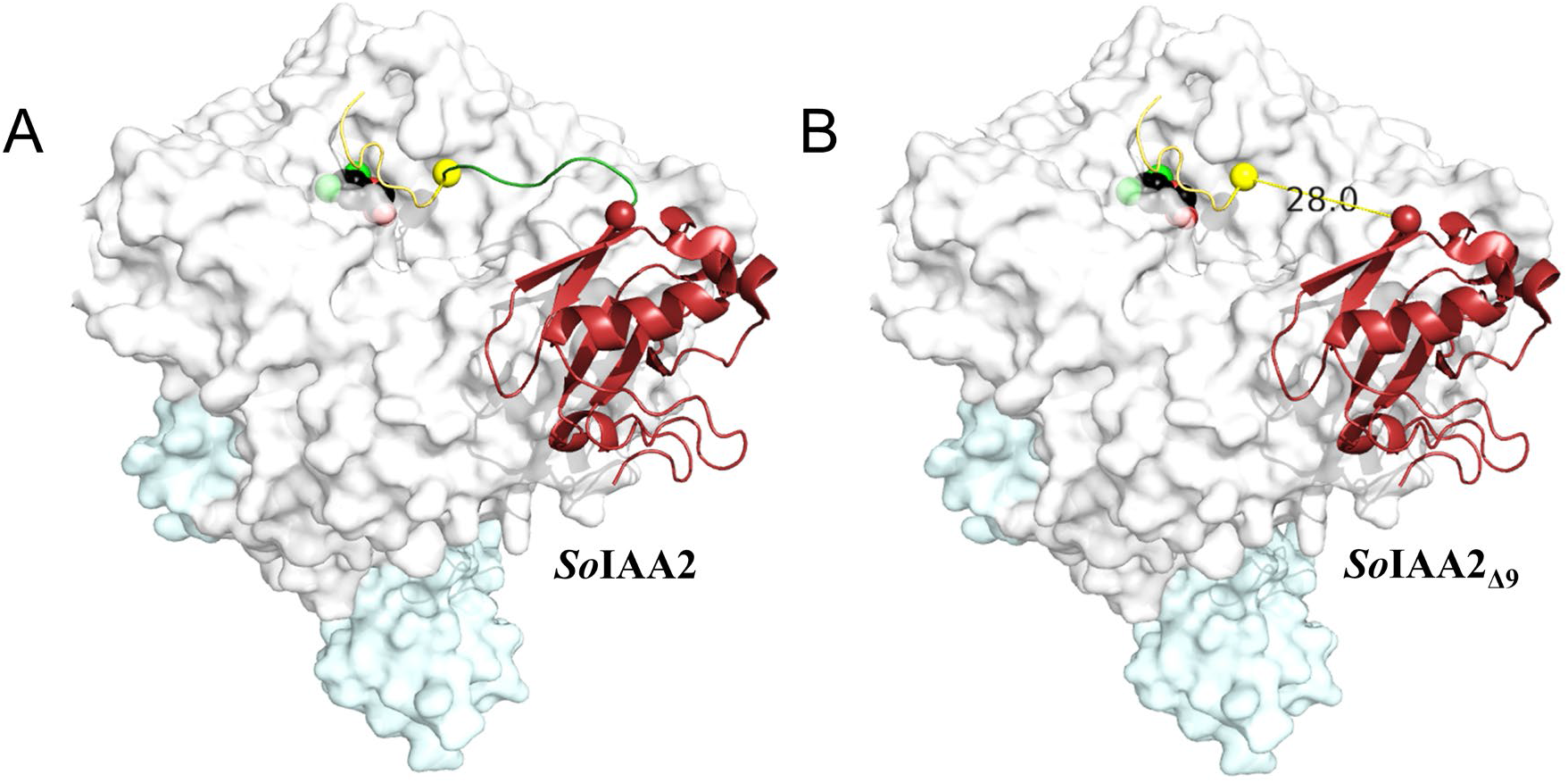
Simulation of the consequences of the IAA2_Δ9_ deletion on the association of the auxin/IAA protein with TIR1/auxin complex. A) The crystallographic model shows ASK1 in light blue, TIR1 in light grey, 2,4-D in black spheres (PDB – 2P1N), and the components of the Aux/IAA protein in three different colors, core degron in yellow (PDB – 2P1N), degron tail in green, and PB1 domain in red (PDB – 2M1M). The TIR1 and Aux/IAA-PB1 interaction residues were manually adjusted in accordance with the HADDOCK simulations performed by Niemeyer et al. (2020). Spheres are used to denominate the region that is lost in IAA2_Δ9_, as seen in (B). The loss of 9 amino acids reduces the distance between domain II and domain III by 28 Angstroms, which is predicted to pull one or both domains away from their binding sites and, hence, reduce the lifetime of the TIR1/auxin/IAA2 auxin complex.

**Figure 5.**
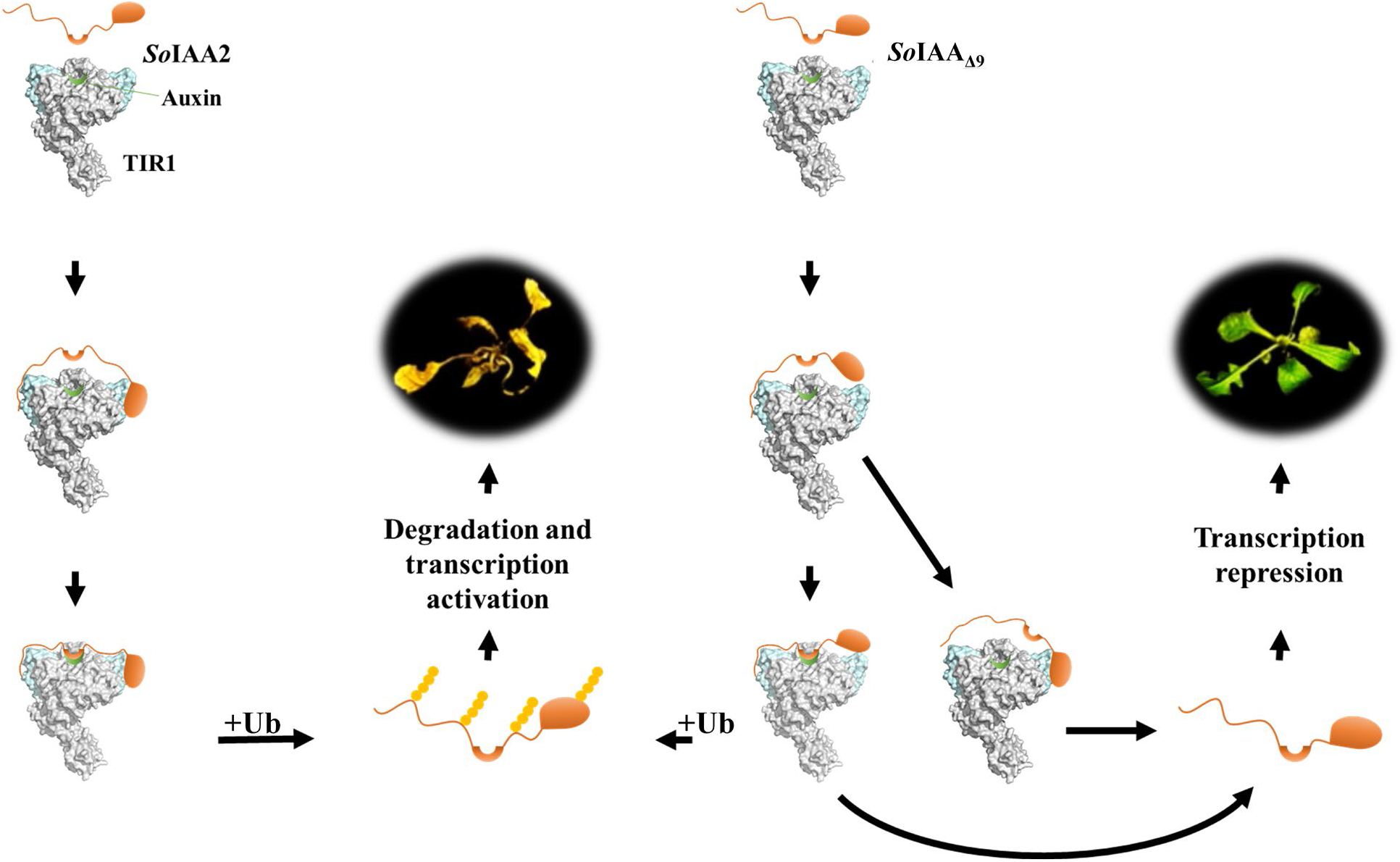
Predicted mechanistic effects of the DT deletion in IAA2_Δ9_ on its association with the TIR1/2,4-D/IAA2 protein complex based on Niemeyer et al. (2020). Left, in the WT *So*IAA2 protein, the degron binds to TIR1 with auxin resulting in ubiquitination and degradation of *So*_IAA2_ by the 26S proteasome, inducing transcriptional activation, which culminates in plant death. Right, in the mutant *So*IAA_2Δ9_ protein, the TIR1 protein may bind the degron but the loss of DT leads to fewer and shorter binding events reducing ubiquitination (based on SPR analysis). In a second possibility, the PB1 domain binds to cluster 1 and prevents the degron from reaching its binding site, reducing binding and consequent ubiquitination (based on the Pymol simulation). Both pathways maintain higher levels of *So*IAA2_Δ9_ in the cell and the associated transcriptional repression of auxin-regulated genes, conferring resistance in the presence of auxin herbicides.

## DISCUSSION

The deletion of 27 nucleotides in the gene *SoIAA2* resulting in the loss of nine amino acids in the DT of *So*IAA2 is a novel mechanism of target site mutation conferring resistance to auxin herbicides in weeds. Furthermore, our results confirm the importance of the DT as a contributor to Aux/IAA stability (Niemeyer et al., 2020) and for perception of synthetic auxin herbicides *in planta.* One explanation for the loss of sensitivity to auxins is the loss of a crucial lysine within the DT, K74. Equivalent residues in the DT of *At*IAA7 and *At*IAA12 were shown to be ubiquitinated in Arabidopsis, but because additional lysine residues are present in IAA2 DT (Niemeyer et al., 2020), the loss of K74 alone is not likely to explain the phenotype. Therefore, we tested the binding interaction of *So*IAA2 with the receptors TIR1 and AFB5 using SPR (Fig. 3), finding that the loss of the DT reduced the affinity of the mutant Aux/IAA protein for both classes of receptor with all auxins tested. A protein model using pre-existing crystallographic structures of TIR1 and the PB1 domain showed that the loss of 9 amino acids in the DT would prevent the simultaneous binding of the core degron and the PB1 domain. Thus, resistance is associated with reduced binding of the mutant *So*IAA2_Δ9_ causing reduced ubiquitination and degradation by the proteasome. The *So*IAA2_Δ9_ protein thus remains in the cell as a transcriptional repressor of ARFs in the presence of synthetic auxin herbicides, preventing the rapid and lethal increase in the gene expression of auxin responsive genes that occurs in susceptible plants when the WT *So*IAA2 is rapidly degraded following auxin herbicide application. *So*IAA2 DT deletion leads to cross-resistance to both 2,4-D and dicamba, revealing exciting options to explore introduction of new synthetic auxin resistance traits in crop plants by targeting the DT region using gene editing technologies.

*SoIAA2* had low expression under natural conditions; however, it was induced rapidly after exogenous auxin application. A similar response was also observed in Arabidopsis Col-0 seedlings, where the expression of *AtIAA2* increased 6 fold one hour after 2,4-D treatment (Yang et al., 2004). The *SoIAA2*_Δ27_ allele encoding the SoIAA2_Δ9_ protein maintained transcriptional repression of auxin responsive genes *GH3.3* and *IAA19* following 2,4-D treatment, showing that the DT is critical for IAA and synthetic auxin herbicide binding leading to Aux/IAA protein degradation. *So*IAA2 seems to play an important role in auxin perception. Previous reports of mutations that confer higher stability to Aux/IAA proteins have been associated with drastic morphological defects, as for *axr5-1/IAA1* (Nagpal et al., 2000; Yang et al., 2015), shy2/IAA3 (Tian & Reed, 1999), *axr2/IAA7* (Nagpal et al., 2000)*, iaa16* (Rinaldi et al., 2012) and *iaa28* (Rogg et al., 2001). All these mutants were reported to be insensitive or hypersensitive to natural and synthetic auxins. Further, in the weed species *Bassia scoparia,* a mutation in the core degron of *BsIAA16* conferred dicamba resistance, but also resulted in a substantial fitness penalty in the form of reduced growth and competitiveness (LeClere et al., 2018; Wu et al., 2021). In our study, over-expression of *SoIAA2_Δ27_* in Arabidopsis induced strong phenotypes in homozygous plants with severely reduced growth (Fig. 2A). Future research is needed to determine whether the *SoIAA2_Δ27_* mutation in 2,4-D resistant *S. orientale* causes any morphological or reproductive defects or reduction in competitiveness with crops.

## MATERIALS AND METHODS

### Plant material

One population of 2,4-D-resistant *S. orientale* was collected from a wheat field in Port Broughton (PB-R), a second 2,4-D resistant population was collected 9.2 km away (PB-R2), while the susceptible population originated from Roseworthy (S), Australia (Preston et al., 2013; Preston & Malone, 2015). Two biparental crosses were made to study inheritance of 2,4-D resistance (Dang et al., 2018), using PB-R as the male parent for Cross A and PB-R2 as the male parent for Cross B, with S as the female parent for both crosses. The F1 progeny were self-pollinated to produce F2 progeny, and F2 individuals were self-pollinated to produce F3 progeny (Preston & Malone, 2015; Dang et al., 2018). Homozygous resistant and homozygous susceptible F3 lines were inbred via single-seed descent to create recombinant inbred lines (RILs) to the F4 generation (Cross A). The F3 generation was used for Cross B.

### RNA-Seq

RNA-Seq was performed on untreated plants from a total of twelve individual RILs, including six individuals each from Cross A (three homozygous 2,4-D-resistant and three homozygous 2,4-D-susceptible) and Cross B (two homozygous 2,4-D-resistant and four homozygous 2,4-D-susceptible). RNA from the youngest expanded leaf of four-leaf stage plants was extracted using the RNeasy Plant Mini Kit (Qiagen) and libraries were prepared using the TruSeq Stranded mRNA Sample Prep Kit (Illumina). Sequencing was performed on a HiSeq 4000 (Illumina) using bar-coded adapters in two lanes of the Illumina flow-cell for 101 cycles, yielding 1.6 billion paired-end reads. Individual library yields ranged from 52 to 75 million paired-end reads. Fastq files were generated and demultiplexed with bcl2fastq v2.17.1.13 conversion software (Illumina) and adaptors were trimmed from the 3’-end reads. The sequence data and processed data files discussed in this publication are deposited in NCBI’s Gene Expression Omnibus (Edgar et al., 2002) and accessible through GEO Series accession number GSE159202 (https://www.ncbi.nlm.nih.gov/geo/query/acc.cgi?acc=GSE159202).

A *de novo* reference transcriptome was obtained by separately assembling Illumina reads from a Cross A 2,4-D-resistant RIL (R206) and a susceptible RIL (S242) with Trinity (Li & Godzik, 2006). The assembled contigs were filtered for minimum contig length of 500 bp. The two assemblies were compared and merged using CDHit v4.6.6 (Li & Godzik, 2006) with a 95% confidence interval to eliminate redundant contigs and retain all contigs unique to either the R or S assembly. This resulted in a 72-Mb transcriptome with 40,987 contigs (available at GEO Series accession number GSE159202) (BUSCO score of 98.4%, Eukaryota ODB10). Putative annotations were assigned using Trinotate (Haas et al., 2013) and the TAIR10 protein database (Berardini et al., 2015). Read alignments to the *de novo* reference transcriptome were conducted with Bowtie2 (Langmead & Salzberg, 2012) using the default “end-to-end” mode and the “sensitive” option. The minimum allowed fragment length was set to 100 and the maximum to 800 bp.

### Differential expression and SNP analysis

Raw read counts were extracted using SAMtools (Li et al., 2009). Counts-per-million (CPM) and gene expression differences were calculated with the package ‘edgeR’ (Robinson et al., 2010) using the statistical software R v3.3 (Team, 2013) and an expression threshold of ≥1 CPM in at least two samples. After normalization expression differences were compared between 2,4-D resistant and susceptible RILs within Cross A and B, respectively, as well as between all 2,4-D-resistant and -susceptible RILs. Differentially expressed transcripts were then filtered for a fold-change of ≥ |2| and a false discovery rate (FDR) adjusted p-value ≤ 0.05. Transcripts annotated as PRP39-2 (10 fold lower in R than S, FDR<0.0001, similar to AT5G46400 that encodes for tetratricopeptide repeat (TPR-like) superfamily protein) and ABCB13 (6 fold higher in R than S, FDR <0.01, similar to AT1G27940 that encodes an auxin efflux transporter) met the criteria for significant differential expression, but were excluded from the resistance hypothesis because both PRP39-2 and ABCB13 had at least one 2,4-D resistant RIL replicate that deviated in expression pattern from the other replicates. Single nucleotide polymorphism (SNP) calling was performed using SAMtools (v. 1.3.1) with the default options and the “mpileup” command. The command “bcftools” was used to retain only SNPs that had a quality score higher than 10 and read depth higher than 10. An additional filtering step was performed for SNPs that were heterozygous or homozygous in at least three individuals of all R-RILs and all S-RILs. Contigs with annotations in the TIR1/AFB family and the Aux/IAA gene family were manually inspected for sequence variants between R-RILs and S-RILs using IGV to view BAM file read alignments (Thorvaldsdóttir et al., 2013).

### Sequence verification of *IAA2* deletion, segregation and analysis of field populations

The *SoIAA2* deleted region was sequenced from samples of R-RILs and S-RILs used in RNAseq. cDNA was synthesized using Iscript (Bio-Rad) and a PCR was performed with primers (Supplementary Table 1) spanning the predicted deletion in *IAA2* and amplicons was determined by Sanger sequencing. To analyze the *IAA2* deletion in other resistant *S. orientale* populations collected in South Australia, plants were sprayed with 200 g ai ha^−1^ 2,4-D at the five true leaf stage using an overhead track sprayer (DeVries Manufacturing) equipped with a flat-fan nozzle tip (Teejet^®^ 8002EVS, Spraying System) calibrated to deliver 187 L ha^−1^ of spray solution at 172 kPa. Percent damage was evaluated at 28 DAT and DNA was extracted by CTAB (Doyle & Doyle, 1987) and analyzed using the same set of primers described above.

A progeny test segregation analysis was performed on 219 F3 plants derived from self-pollination of a heterozygous F2 individual from cross B, identified via progeny test due to segregation of 2,4-D resistance in the F3. Plants were sprayed with 250 g ai ha^−1^ 2,4-D at the five true leaf stage as described above. KASP genotyping markers were developed using a forward primer specific to the R (HEX) and S (FAM), together with a universal reverse primer (Supplementary Table 1).

### Functional validation of *IAA2* deletion in Arabidopsis

*SoIAA2* alleles were amplified by PCR from cDNA (Supplemental Table 1), ligated into the binary vector *pFGC5941* and transformed into *Agrobacterium tumefaciens* (strain GV3101). Each allele of So*IAA2* was transformed into *Arabidopsis thaliana* (Col-0) by floral dip method (Clough & Bent, 1998). Seeds were plated on ½ MS, 1% phytoagar, and 7.5 μg mL^−1^ of ammonium glufosinate for selection of transformed seedlings. Homozygous plants of the subsequent generations containing one copy of the transgene were selected.

Root assays were performed in ½ MS and 1% phytoagar supplemented with synthetic and natural auxins (2,4-D, dicamba, and IAA, Caisson Labs). Seeds of Arabidopsis were gas sterilized, incubated at 4°C in dark, plated directly in media with the respective treatment or DMSO as a control and moved to a growth chamber after 3 days (60% RH, 21/18 °C, 16/8 h light/dark photoperiod). Roots were photographed and measured using ImageJ (Abràmoff et al., 2004) 14 d after the plates were moved to the growth conditions. Post emergence dose responses with 2,4-D and dicamba were performed on pre-flowering 21 d old homozygous transgenic plants for all constructs. Plants were harvested 21 d after treatment, dry mass was measured, and the dataset was processed using the R package drc (Knezevic et al., 2007) to generate dose response models (Supplementary Table 2).

For gene expression analysis under auxin treatments, soil cultivated *S. orientale* (5-6 leaves) and transgenic Arabidopsis lines (pre-flowering 16 leaves) were sprayed with 250 g ai ha^−1^ 2,4-D and at 6 h after treatment a leaf disc was collected from the 8^th^ leaf in Arabidopsis and 4^th^ for *S. orientale,* cDNA was generated, and gene expression of known Arabidopsis auxin-responsive genes *GH3.3; AtIAA19* and *AtIAA2* was analyzed by qPCR (Supplementary Table 1).

### *In silico* protein structure predictions

Susceptible (*So*IAA2) and resistant (*So*IAA2_Δ9_) protein sequences were used for IDR predictions using PrDos (Ishida & Kinoshita, 2007), IUPred (Mészáros et al., 2018) and Spot2 (Hanson et al., 2016) algorithms. Kyte-Doolittle hydropathy maps were calculated by Expasy-linked ProtScale (Gasteiger et al., 2005).

### IAA2 protein purification

The cDNAs corresponding to the two alleles of *SoIAA2* were cloned into a pFN2A (GST) flexi vector (Promega – primers in Supplementary Table 1), expressed in *E. coli* BL21 and purified using GST beads (Glutathione Sepharose 4B from GE), according to the manufacturer manual. The GST-IAA2 proteins were recovered, GST was cleaved using TEV protease (100 μL at 400 μM – New England Biolabs), and GST beads were added to eliminate GST and undigested proteins. The final purified *So*IAA2 and *So*IAA2_Δ9_ proteins were then quantified using spectrometry and separated on SDS-PAGE to verify the purity.

### SEC purification and retention volume studies

The cleaved IAA2 proteins were loaded into a GE Superdex 200 10/300 increase column equilibrated in CD buffer (10 mM Tris H_2_SO_4_ pH 7.0 and 150 mM NaClO_4_) and quantified using a Cary Bio-50 UV-Vis spectrophotometer. Proteins were quantified by a UV 280 nm detector. Protein oligomers were analyzed based on their retention volumes and compared to Bio-Rad gel filtration protein standards (15 – 600 kDa).

### Circular dichroism (CD) analysis

CD was performed using a MOS-500 spectrometer (Bio-logic, France) scanning 185-265 nm with a 2 nm slit width at an acquisition rate of 2 sec nm^−1^. Buffer and proteins were scanned in a PTFE stoppered 1 mm UV quarts cuvette (FireflySci). Protein concentration was between 1 to 7 μM, and the absorbance and PMT voltage targeted were 1 AUand 500 V, respectively. CD measurements were taken three times and the raw ellipticity values were averaged after buffer subtraction and converted to MRE or μMol ellipticity values.

### Surface Plasmon Resonance (SPR) affinity binding analysis

SPR experiments were conducted according to the protocols described in Lee et al. (2014) with modifications. TIR1/AFB5 was expressed in Sf9 insect cell culture using a recombinant baculovirus encoding 10xHis:GFP:FLAG:TEV:receptor and 10xHis:ASK1. Initial purification using the His-tag on nickel resin was followed by clean-up using FLAG chromatography. For SPR assays using a Biacore T200 (Cytiva), the purified protein was mixed with the appropriate concentration of each auxin before passing it over a streptavidin chip loaded with biotinylated *So*IAA2 and *So*IAA2_Δ9_ degron peptides (Thermo Fisher Scientific) (Supplementary Table 3).

The SPR buffer was Hepes-buffered saline (10 mM Hepes, 3 mM EDTA, 150 mM NaCl and 0.05% Tween 20). Binding experiments were run at a flow rate of 30 μl min^−1^ using a series of 100 s of injection and dissociation times, followed by a final dissociation of 600 s, according to the manufacturer’s single cycle kinetic routine. Data from a control channel (loaded with a mutated degron, mIAA7) (Lee et al., 2014) and from a buffer+DMSO-only run were subtracted from each sensogram following the standard double reference subtraction protocol.

### Crystallographic models Pymol for manual docking of *So*IAA2 and *So*IAA2_Δ9_ and TIR1 association

A crystallographic model of the co-receptor complex was simulated based on the available model of Arabidopsis ASK1-TIR1 in association with 2,4-D and a 13 amino acid degron motif (PDB: 2P1N) and the PB1 domain structure from pea (*Pisum sativum*, PDB: 2M1M). The PB1 domain was manually adjusted to give the interactions simulated using molecular dynamics by Niemeyer et al. (2020) using their model for *At*IAA7 on *At*TIR1. For TIR1, we used D119, D170, V171, S172, H174, H178, S199, and R220; and for PB1 of Aux/IAA, we used Y196, R225, P237, and R238. The molecular distance corresponding to the length of the missing DT was measured based on the position of the C-terminal amino-acid of the core degron when bound (PDB: 2P1N) and the first amino acid of the PB1 domain positioned on its binding cluster (after Niemeyer et al., 2020).

## ACKNOWLEDGEMENTS

The authors thank Rachel Chayer for assistance with lab work. This work was supported in part by funding from Dow AgroSciences, by the Grains Research and Development Corporation – Australian Government, grant UA00158, the Brazilian governmental scholarship from the National Council for Scientific and Technological Development (CNPq – 207387/2014-1) to M.F., and by the EU MSCA-IF project CrysPINs (792329) in support of MK.

## Notes

### Competing Interest Statement

The authors have declared no competing interest.

## REFERENCES

Abràmoff M.D., Magalhães, P.J. and Ram, S.J. (2004) Image processing with ImageJ. Biophotonics Intl. 11, 36–42.

Behrens M.R., Mutlu, N., Chakraborty, S., Dumitru, R., Jiang, W.Z., LaVallee, B.J., Herman, P.L., Clemente, T.E. and Weeks, D.P. (2007) Dicamba resistance: Enlarging and preserving biotechnology-based weed management strategies. Science 316, 1185–1188.

Berardini T.Z., Reiser, L., Li, D., Mezheritsky, Y., Muller, R., Strait, E. and Huala, E. (2015) The Arabidopsis information resource: Making and mining the “gold standard” annotated reference plant genome. Genesis 53, 474–485.

Calderón Villalobos L.I.A., Lee, S., De Oliveira, C., Ivetac, A., Brandt, W., Armitage, L., Sheard, L.B., Tan, X., Parry, G., Mao, H., Zheng, N., Napier, R., Kepinski, S. and Estelle, M. (2012) A combinatorial TIR1/AFB-Aux/IAA co-receptor system for differential sensing of auxin. Nat. Chem. Biol. 8, 477–485.

Clough S.J. and Bent, A.F. (1998) Floral dip: a simplified method for *Agrobacterium-mediated* transformation of *Arabidopsis thaliana*. Plant J. 16, 735–743.

Dang H.T., Malone, J.M., Boutsalis, P., Krishnan, M., Gill, G. and Preston, C. (2018) Reduced translocation in 2, 4-D-resistant oriental mustard populations *(Sisymbrium orientale* L.) from Australia. Pest Manage. Sci. 74, 1524–1532.

Dinesh D.C., Kovermann, M., Gopalswamy, M., Hellmuth, A., Calderón Villalobos, L.I.A., Lilie, H., Balbach, J. and Abel, S. (2015) Solution structure of the *Ps*IAA4 oligomerization domain reveals interaction modes for transcription factors in early auxin response. Proc. Natl. Acad. Sci. USA 112, 6230–6235.

Dinesh D.C., Villalobos, L.I.A.C. and Abel, S. (2016) Structural biology of nuclear auxin action. Trends Plant Sci. 21, 302–316.

Doyle J.J. and Doyle, J.L. (1987) A rapid DNA isolation procedure for small quantities of fresh leaf tissue. Phytochemical Bulletin 19, 11–15.

Edgar R., Domrachev, M. and Lash, A.E. (2002) Gene Expression Omnibus: NCBI gene expression and hybridization array data repository. Nuc. Acids Res. 30, 207–210.

Figueiredo M.R., Leibhart, L.J., Reicher, Z.J., Tranel, P.J., Nissen, S.J., Westra, P., Bernards, M.L., Kruger, G.R., Gaines, T.A. and Jugulam, M. (2018) Metabolism of 2,4-dichlorophenoxyacetic acid contributes to resistance in a common waterhemp (*Amaranthus tuberculatus*) population. Pest Manage. Sci. 74, 2356–2362.

Gaines T.A., Duke, S.O., Morran, S., Rigon, C.A.G., Tranel, P.J., Küpper, A. and Dayan, F.E. (2020) Mechanisms of evolved herbicide resistance. J. Biol. Chem. 295, 10307–10330.

Gasteiger E., Hoogland, C., Gattiker, A., Duvaud, S.e., Wilkins, M.R., Appel, R.D. and Bairoch, A. 2005. Protein identification and analysis tools on the ExPASy server. In: Walker JM ed. The Proteomics Protocols Handbook. Totowa, NJ: Humana Press, 571–607.

Goggin D.E., Bringans, S., Ito, J. and Powles, S.B. (2020) Plasma membrane receptor-like kinases and transporters are associated with 2,4-D resistance in wild radish. Ann. Bot. 125, 821–832.

Goggin D.E., Cawthray, G.R. and Powles, S.B. (2016) 2,4-D resistance in wild radish: reduced herbicide translocation via inhibition of cellular transport. J. Exp. Bot. 67, 3223–3235.

Haas B.J., Papanicolaou, A., Yassour, M., Grabherr, M., Blood, P.D., Bowden, J., Couger, M.B., Eccles, D., Li, B. and Lieber, M. (2013) *De novo* transcript sequence reconstruction from RNA-seq using the Trinity platform for reference generation and analysis. Nat. Protoc. 8, 1494.

Hanson J., Yang, Y., Paliwal, K. and Zhou, Y. (2016) Improving protein disorder prediction by deep bidirectional long short-term memory recurrent neural networks. Bioinformatics 33, 685–692.

Heap I. (2020) The international survey of herbicide resistant weeds. Available on-line: https://www.weedscience.com. Accessed April 22, 2020.

Ishida T. and Kinoshita, K. (2007) PrDOS: prediction of disordered protein regions from amino acid sequence. Nuc. Acids Res. 35, W460–W464.

Kim J., Harter, K. and Theologis, A. (1997) Protein-protein interactions among the Aux/IAA proteins. Proc. Natl. Acad. Sci. USA 94, 11786–11791.

Knezevic S.Z., Streibig, J.C. and Ritz, C. (2007) Utilizing R software package for dose-response studies: The concept and data analysis. Weed Technol. 21, 840–848.

Langmead B. and Salzberg, S.L. (2012) Fast gapped-read alignment with Bowtie 2. Nat. Methods 9, 357–359.

LeClere S., Wu, C., Westra, P. and Sammons, R.D. (2018) Cross-resistance to dicamba, 2,4-D, and fluroxypyr in *Kochia scoparia* is endowed by a mutation in an *AUX/IAA* gene. Proc. Natl. Acad. Sci. USA 115, E2911–E2920.

Lee S., Sundaram, S., Armitage, L., Evans, J.P., Hawkes, T., Kepinski, S., Ferro, N. and Napier, R.M. (2014) Defining binding efficiency and specificity of auxins for SCF^TIR1/AFB^-Aux/IAA co-receptor complex formation. ACS Chem. Biol. 9, 673–682.

Li H., Handsaker, B., Wysoker, A., Fennell, T., Ruan, J., Homer, N., Marth, G., Abecasis, G. and Durbin, R. (2009) The sequence alignment/map format and SAMtools. Bioinformatics 25, 2078–2079.

Li W. and Godzik, A. (2006) Cd-hit: a fast program for clustering and comparing large sets of protein or nucleotide sequences. Bioinformatics 22, 1658–1659.

Mészáros B., Erdős, G. and Dosztányi, Z. (2018) IUPred2A: context-dependent prediction of protein disorder as a function of redox state and protein binding. Nuc. Acids Res. 46, W329–W337.

Micsonai A., Wien, F., Bulyáki, É., Kun, J., Moussong, É., Lee, Y.-H., Goto, Y., Réfrégiers, M. and Kardos, J. (2018) BeStSel: a web server for accurate protein secondary structure prediction and fold recognition from the circular dichroism spectra. Nuc. Acids Res. 46, W315–W322.

Mockaitis K. and Estelle, M. (2008) Auxin receptors and plant development: a new signaling paradigm. Ann. Rev. Cell Devel. Biol. 24, 55–80.

Nagpal P., Walker, L.M., Young, J.C., Sonawala, A., Timpte, C., Estelle, M. and Reed, J.W. (2000) *AXR2* encodes a member of the Aux/IAA protein family. Plant Physiol. 123, 563–574.

Nanao M.H., Vinos-Poyo, T., Brunoud, G., Thévenon, E., Mazzoleni, M., Mast, D., Lainé, S., Wang, S., Hagen, G., Li, H., Guilfoyle, T.J., Parcy, F., Vernoux, T. and Dumas, R. (2014) Structural basis for oligomerization of auxin transcriptional regulators. Nature Comm, 5, 3617.

Niemeyer M., Moreno Castillo, E., Ihling, C.H., Iacobucci, C., Wilde, V., Hellmuth, A., Hoehenwarter, W., Samodelov, S.L., Zurbriggen, M.D., Kastritis, P.L., Sinz, A. and Calderón Villalobos, L.I.A. (2020) Flexibility of intrinsically disordered degrons in AUX/IAA proteins reinforces auxin co-receptor assemblies. Nature Comm, 11, 2277.

Pettinga D.J., Ou, J., Patterson, E.L., Jugulam, M., Westra, P. and Gaines, T.A. (2018) Increased Chalcone Synthase (CHS) expression is associated with dicamba resistance in *Kochia scoparia*. Pest Manage. Sci. 74, 2306–2315.

Preston C., Dolman, F.C. and Boutsalis, P. (2013) Multiple resistance to acetohydroxyacid synthase-inhibiting and auxinic herbicides in a population of oriental mustard *(Sisymbrium orientale)*. Weed Sci. 61, 185–192.

Preston C. and Malone, J.M. (2015) Inheritance of resistance to 2, 4-D and chlorsulfuron in a multiple resistant population of *Sisymbrium orientale*. Pest Manage. Sci. 71, 1523–1528.

Rinaldi M.A., Liu, J., Enders, T.A., Bartel, B. and Strader, L.C. (2012) A gain-of-function mutation in IAA16 confers reduced responses to auxin and abscisic acid and impedes plant growth and fertility. Plant Mol. Biol. 79, 359–373.

Robinson M.D., McCarthy, D.J. and Smyth, G.K. (2010) edgeR: a Bioconductor package for differential expression analysis of digital gene expression data. Bioinformatics 26, 139–140.

Rogg L.E., Lasswell, J. and Bartel, B. (2001) A gain-of-function mutation in *IAA28* suppresses lateral root development. Plant Cell 13, 465–480.

Tan X., Calderon-Villalobos, L.I.A., Sharon, M., Zheng, C., Robinson, C.V., Estelle, M. and Zheng, N. (2007) Mechanism of auxin perception by the TIR1 ubiquitin ligase. Nature 446, 640–645.

Tao S. and Estelle, M. (2018) Mutational studies of the Aux/IAA proteins in *Physcomitrella* reveal novel insights into their function. New Phytol. 218, 1534–1542.

Team R.C. (2013) R: A language and environment for statistical computing.

Thorvaldsdóttir H., Robinson, J.T. and Mesirov, J.P. (2013) Integrative Genomics Viewer (IGV): high-performance genomics data visualization and exploration. Brief. Bioinform. 14, 178–192.

Tian Q. and Reed, J.W. (1999) Control of auxin-regulated root development by the *Arabidopsis thaliana SHY2/IAA3* gene. Development 126, 711–721.

Todd O.E., Figueiredo, M.R.A., Morran, S., Soni, N., Preston, C., Kubeš, M.F., Napier, R. and Gaines, T.A. (2020) Synthetic auxin herbicides: finding the lock and key to weed resistance. Plant Sci. 300, 110631.

Ulmasov T., Liu, Z.B., Hagen, G. and Guilfoyle, T.J. (1995) Composite structure of auxin response elements. Plant Cell 7, 1611–1623.

Wright T.R., Shan, G., Walsh, T.A., Lira, J.M., Cui, C., Song, P., Zhuang, M., Arnold, N.L., Lin, G. and Yau, K. (2010) Robust crop resistance to broadleaf and grass herbicides provided by aryloxyalkanoate dioxygenase transgenes. Proc. Natl. Acad. Sci. USA 107, 20240–20245.

Wu C., Paciorek, M., Liu, K., LeClere, S., Perez-Jones, A., Westra, P. and Sammons, R.D. (2021) A dicamba resistance endowing IAA16 mutation leads to significant vegetative growth defects and impaired competitiveness in kochia (*Bassia scoparia*). Pest Manage. Sci., in press.

Yang X., Lee, S., So, J.-h., Dharmasiri, S., Dharmasiri, N., Ge, L., Jensen, C., Hangarter, R., Hobbie, L. and Estelle, M. (2004) The IAA1 protein is encoded by AXR5 and is a substrate of SCFTIR1. Plant J. 40, 772–782.

Yang Y., Zhang, X. and Yu, B. (2015) O-Glycosylation methods in the total synthesis of complex natural glycosides. Nat. Prod. Rep. 32, 1331–1355.

