## Supplementary Information for "An in-frame deletion mutation in the degron tail of auxin co-receptor *IAA2* confers resistance to the herbicide 2,4-D in *Sisymbrium orientale*"

^b^ Bayer AG, Division CropScience, Weed Control Research, Building H872, 65926 Frankfurt, Germany.

^c^ School of Agriculture, Food and Wine, University of Adelaide, PMB1 Glen Osmond 5064, South Australia, Australia.

^d^ Department of Biochemistry and Molecular Biology, Colorado State University, 1870 Campus Delivery, Fort Collins, CO 80523, USA.

^e^ Department of Biology and Program in Cell and Molecular Biology, Colorado State University, 1878 Campus Delivery, Fort Collins, CO 80523, USA.

^f^ Department of Plant, Soil, and Microbial Sciences, Michigan State University, East Lansing, MI 48824, USA.

^g^ School of Life Sciences, The University of Warwick, Coventry, CV4 7AL, UK.

*Corresponding Author: Todd A. Gaines. Department of Agricultural Biology, Colorado State University, 1177 Campus Delivery, Fort Collins, CO 80523, USA.. Ph: +1 970 491 6824.

Supplementary Table 1. Primer list.

| **Experiment** | **Primer** | **Sequence** |
| --- | --- | --- |
| **Candidate gene validation** | PCR genotyping | Amplicon size testing primer  FW 5’-AACTCAAATCGTTGGTTGGC-3’  Resistant specific  FW 5’- CGTAAGAACAACAACAGTGTGAGC -3’  Susceptible specific  FW 5’- CTCCGGTGAGATCTTATGTG -3’  Universal reverse  RV 5’-CTTTATCCTCGTACGTTGGTACG-3’, |
|  | KASP genotyping | HEX-FW 5’- GAAGGTCGGAGTCAACGGATT*CTCCGGTGAGATCTTATGTG*-3’  FAM-FW 5’- GAAGGTGACCAAGTTCATGCT*CGTAAGAACAACAACAGTGTGAGC*-3’  Universal RV 5’-ATTTTGCGGAGGTATGGTGC-3’ |
| **Arabidopsis transformation** | pFGC5941 cloning using AscI and BamHI as restriction sites | 1AA2 -AscI-FW 5’-TTGGCGCGCC ATGGCGTACGAGAAAGTCAATGAGCTTA-3’  1AA2- BamHI-RV 5'-CGGGATCCTCATAAGGAAGAGTCATCAGATCCTTTCATGATTC-3' |
| ***IAA2* expression in *Sisymbrium orientale*** | *Aux/IAA19* | FW 5’- GCGTCGTTTCATCAGGTAGT -3’  RV 5’- TTCCTCCGGTAAGAACAAACC -3’ |
|  | *GH3.3* | FW 5’- ATCAAAGACTCCTGGTGGATTAC -3’  RV 5’- ACACGTTGTACGGATCGTATG -3’ |
|  | *SoIAA2*_WT_ | FW 5’- GAACAACAACAGTGTGAGCTATG -3’  RV 5’- GCCTTGAGAAGCTCTGGATAG -3’ |
|  | *SoIAA2*_Δ27_ | FW 5’- CTCCGGTGAGATCTTATGTG -3’  RV 5’- CTCCGGTGAGATCTTATGTG-3’ |
|  | *SoCyclophilin* | FW 5’- CATGTGCCAAGGAGGAGATT-3’  RV 5’-GTGTGCTTCCTCTCGAAGTT-3’ |
|  | *SoActin2* | FW 5’- GTGGAACCACTATGTTCTCTGG-3’  RV 5’- GGAGGTGCAACGACCTTAAT-3’ |
| ***IAA2* expression in *Arabidopsis thaliana*** | *Aux/IAA19* | FW 5’- AGGACTCGGGCTTGAGATAA -3’  RV 5’- CTCCGTGAAAGCTCTCTTCTTC -3’ |
|  | *GH3.3* | FW 5’- GAGAGCAAGGAACTCGTGTTAT -3’  RV 5’- CGTCATTTGGAGGATTGGTTTG -3’ |
|  | *AtCyclophilin* | FW 5’- GTCTGATAGAGATCTCACGT -3’  RV 5’- AATCGGCAACAACCACAGGC -3’ |
|  | *AtActin2* | FW 5’- GGCAAGTCATCACGATTGG -3’  RV 5’- CAGCTTCCATTCCCACAAAC -3’ |
| **IAA2 expression in *E. coli*** | pFN2A (GST) Flexi Vector cloning using SgfI and PmeI as restriction sites | GST-IAA2-SgfI-FW 5’- ttGCGATCGCGGCGTACGAGAAAGTCAATG-3’  GST-IAA2-PmeI-RV 5’-ttGTTTAAACTCATAAGGAAGAGTCATCAGATCCTTTCATGATTC-3’ |

**Supplementary Table 2.** Equation parameters for the graphs to fit the dose response data of transgenic lines of Arabidopsis (graphs shown in Supplementary Figure 4).

| **Graph** | **Line** | **Equation** |
| --- | --- | --- |
| **2,4-D** | IAA2 R1 | f(x) = 0 + ((101.93 – 0 + 0.34x) / (1 + exp(2.15(log(x) − log(664.34))))) |
|  | IAA2 R4 | f(x) = 0 + ((99.59 – 0 + 10.43x) / (1 + exp(1.19(log(x) − log(8.82))))) |
|  | IAA2 S7 | f(x) = 7.5834 + 7015 − 7.5834 (1 + exp(0.096489 (x – (-44.725)))) |
|  | IAA2 S1 | f(x) = 11.682 + 7273.8 − 11.682 (1 + exp(0.022167 (x – (-199.94)))) |
|  | Ø 4 | f(x) = 10.904 + 3198.4 − 10.904 (1 + exp(0.045309 (x – (-78.401)))) |
|  | Ø 2 | f(x) = 12.742 + 7223.2 −12.742 (1 + exp(0.031865 (x – (-138.45)))) |
|  | Col-0 | f(x) = 17.321 + 646.53 − 17.321 (1 + exp(0.10772 (x – (-17.533)))) |
| **Dicamba** | IAA2 R1 | f(x) = 0 + ((419.52 – 0 + (-0.12x)) / (1 + exp(-0.51(log(x) − log(405.38))))) |
|  | IAA2 R4 | f(x) = 0 + ((97.56 – 0 + (1.77x)) / (1 + exp(1.36(log(x) − log(115.31))))) |
|  | IAA2 S7 | f(x) = 17.061 + 2554.3 − 17.061 (1 + exp(0.03138 (x – (-107.95)))) |
|  | IAA2 S1 | f(x) = 26.54 + 1207.7 − 26.54 (1 + exp(0.012065 (x – (-225.08)))) |
|  | Ø 4 | f(x) = 39.286 + 2327.7 − 39.286 (1 + exp(0.015672 (x – (-231.74)))) |
|  | Ø 2 | f(x) = 30.379 + 10000 − 30.379 (1 + exp(0.032403 (x – (-153.1)))) |
|  | Col-0 | f(x) = 30.687 + 2077.6 − 30.687 (1 + exp(0.0063358(x – (-536.49)))) |

**Supplementary Table 3. Biotinylated IAA2 degron peptides for SPR analysis.** The peptides contained an N-terminal biotin, the core degron (GWPPVR), with or without the nine amino acids deleted from the degron tail, and 6 amino acids from the PB1 domain.

| Biot-TKT | Degron | Degron tail | Fraction of PB1 |
| --- | --- | --- | --- |
| IAA2 | QIVGWPPVR | SSRKNNNSV | SYVKVS |
| IAA2_Δ9_ | QIVGWPPVR |  | SYVKVS |

**Supplementary Table 4. The *IAA2*_Δ27_ genotype correlates with the 2,4-D resistant phenotype in several field populations of *Sisymbrium orientale* from South Australia.** All plants of the four susceptible populations were homozygous for the *SoIAA2_WT_* S allele (P15, P31, P49, P50). In resistant populations, all the plants tested for PB-R, PB-R2 and P17 were homozygous for the *IAA2_Δ27_* R allele or heterozygous. Resistant population P28 was homozygous for the WT *IAA2*, suggesting that this population evolved a different mechanism for 2,4-D resistance.

| Population | No. of individuals tested | No. of Individuals with Genotype | | |
| --- | --- | --- | --- | --- |
|  |  | IAA2_WT_ | IAA2_Δ27_ | IAA2_WT_/IAA2_Δ27_ |
| PB-R (R) | 20 |  | 20 |  |
| PB-R2 (R) | 20 |  | 20 |  |
| P15 (S) | 15 | 15 |  |  |
| P17 (R) | 20 |  | 14 | 6 |
| P28 (R) | 20 | 20 |  |  |
| P31 (S) | 5 | 5 |  |  |
| P49 (S) | 5 | 5 |  |  |
| P50 (S) | 5 | 5 |  |  |

**Supplementary Table 5.** **The mutant IAA2_Δ9_ binds to the receptors TIR1 and AFB5 with poorer affinity than the WT IAA2.** Surface plasmon resonance was used to measure the binding of purified receptor protein to each degron peptide using single cycle kinetics. Auxins were mixed with the TIR1 or AFB5 preparation in advance of injection over the biotinylated IAA2 peptides using standard Biacore double baseline subtraction with single cycle kinetic routines.

**Kinetic data for the binding of TIR1 to *So*IAA2 and *So*IAA2_Δ9._**

| **Ligand** | **Auxin** | ***k*_a_ (M^-1^ s^-1^)** | ***k*_d_ (s^-1^)** | **K_D_ (M)** |
| --- | --- | --- | --- | --- |
| *So*IAA2 **_Δ9_** | IAA | 1.23E+04 | 5.01E-04 | 4.07E-08 |
| *So*IAA2 | IAA | 1.83E+04 | 2.08E-04 | 1.14E-08 |
| *So*IAA2 **_Δ9_** | 2,4-D | 1.11E+05 | 3.29E-02 | 2.96E-07 |
| *So*IAA2 | 2,4-D | 3.01E+05 | 4.04E-02 | 1.35E-07 |
| *So*IAA2 **_Δ9_** | dicamba | 3.90E+03 | 2.71E-03 | 6.94E-07 |
| *So*IAA2 | dicamba | 7.71E+03 | 1.93E-03 | 2.50E-07 |

**Kinetic data for the binding of AFB5 to *So*IAA2 and *So*IAA2_Δ9._**

| **Ligand** | **Auxin** | ***k*_a_ (M^-1^ s^-1^)** | ***k*_d_ (s^-1^)** | **K_D_ (M)** |
| --- | --- | --- | --- | --- |
| SoIAA2 **_Δ9_** | IAA | 4.35E02 | 1.10E-3 | 2.52E-06 |
| SoIAA2 | IAA | 9.19E02 | 3.26E-3 | 3.54E-06 |
| SoIAA2 **_Δ9_** | 2,4-D | No fit |  |  |
| SoIAA2 | 2,4-D | No fit |  |  |

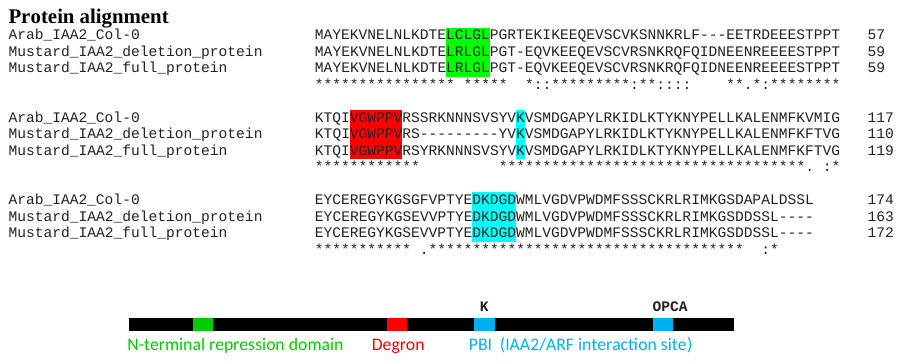

**Supplementary Figure 1. Amino acid alignment of *At*IAA2, *So*IAA2_Δ9_, and *So*IAA2.** The protein alignment of *At*IAA2 with *So*IAA2 shows 87.1% aa identity and 89.9% similarity. The colored residues refer to the conserved elements in all functional domains within Aux/IAA proteins. The ethylene response factor associated amphiphilic repression (EAR) motifs shown in green (amino acid sequence LxLxL) binds to TOPLESS (TPL), a transcriptional corepressor. The sequence highlighted in red corresponds to the degron motif, which binds to auxin and the co-receptor TIR/AFB proteins. In blue, there are the K and DxD and ExD sequences (OPCA) motifs in the PB1 domain. Those two motifs form complementary interaction centers, where K makes one portion of the protein basic and OPCA makes the other portion acidic, leading to the formation of oligomers between Auxin Response Factors (ARF) and other Aux/IAA proteins that have the same highly conserved PB1 domain (Tao & Estelle, 2018). The 9 aa deletion (positions 72-80) in the herbicide resistant populations of *Sisymbrium orientale* occurs between the degron and PB1 domain, a region named the degron tail (Niemeyer et al., 2020).

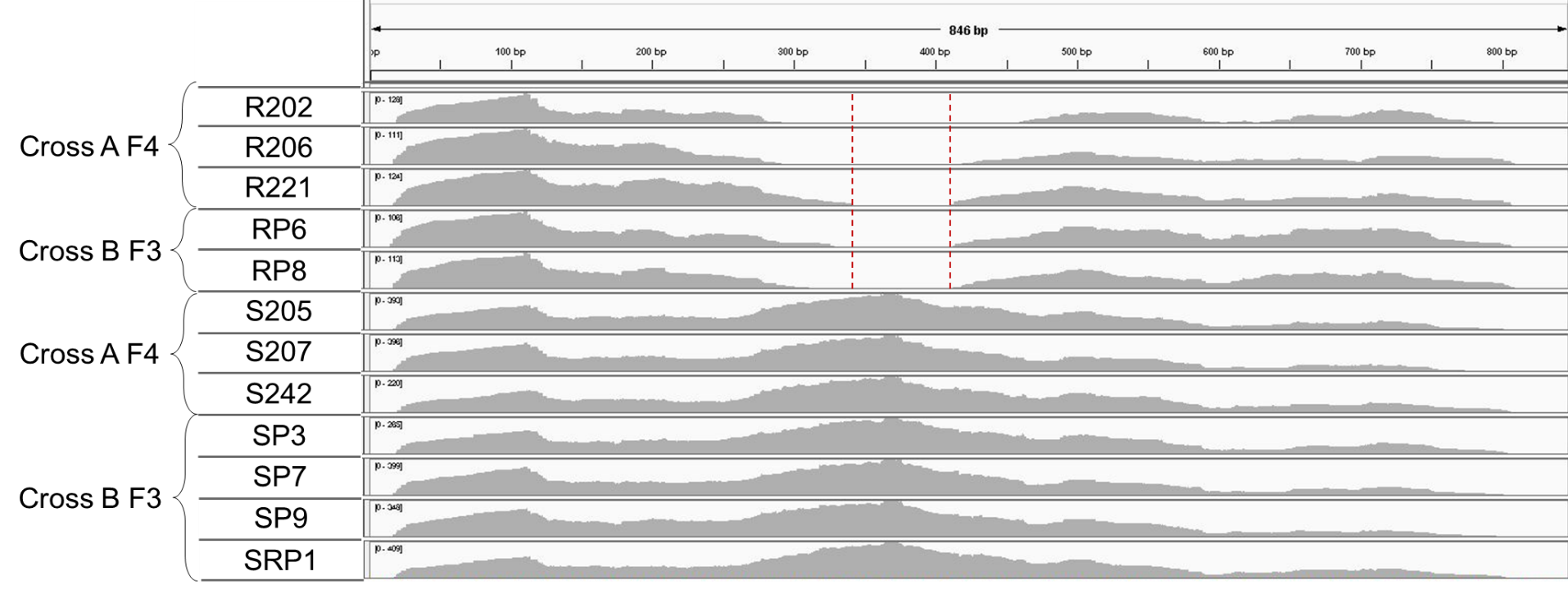

**Supplementary Figure 2. RNA-Seq read depth and PCR assay of 27 bp deletion for confirmation of deletion in *SoIAA2*.** Illumina RNA-Seq read depth of *SoIAA2* transcripts in resistant Recombinant Inbred Lines (RILs), 3 plants from F4 cross A (R 202, 206 and 221) and 2 plants from F3 cross B (RP6 and 8). Followed by susceptible RILs, 3 plants from F4 cross A (S 205, 207 and 242) and 4 plants from F3 cross B (SP3, 7, 9 and SRP1). The identified region with red dotted lines in the resistant RILs shows the 27 bp deletion in the Degron tail (DT) region of *SoIAA2*.

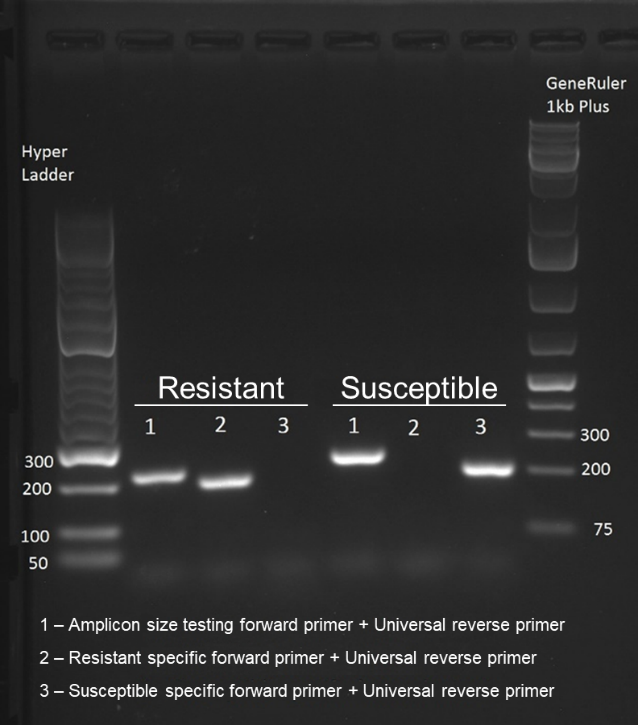
The electrophoresis gel confirms the absence of these 27 bp in resistant plants. Band 1 shows the different sizes between resistant (212 bp) and susceptible (239 bp) *SoIAA2* alleles. The set of primers that generated band 2 corresponds to specific nucleotides flanking the DT deleted region, which amplify only in the resistant allele (band present in resistant but not in susceptible sample). The set of primers for band 3 only anneal to the nucleotides in the DT that are only present in the original wild type *SoIAA2* allele, but this primer set does not amplify in resistant plants (band present in susceptible but absent in resistant sample). The PCR reactions were performed using cDNA synthesized from RNA of R221 and S205 F4 RILs.

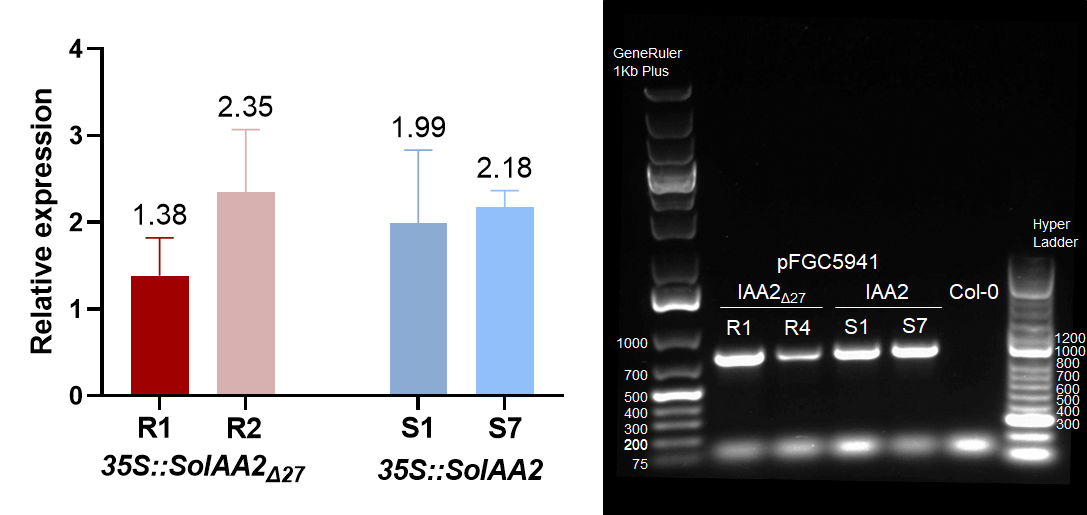

**Supplementary Figure 3. RT-PCR data and confirmation genomic insertion of pFGC5941 T-DNA in Arabidopsis *SoIAA2_Δ27_* and *SoIAA2* transgenic lines.** Relative expression of *SoIAA2* compared to the reference genes *AtCyclophilin* and *AtActin2* in T3 homozygous Arabidopsis lines with a single copy of T-DNA vector. Histograms correspond to the average of three plants of each line with standard error bars. Electrophoresis gel confirmed the genomic insertion of pFGC5941 T-DNA in Arabidopsis transgenic lines, generating amplicons between 800-1000 bp and no amplification for control (Col-0).

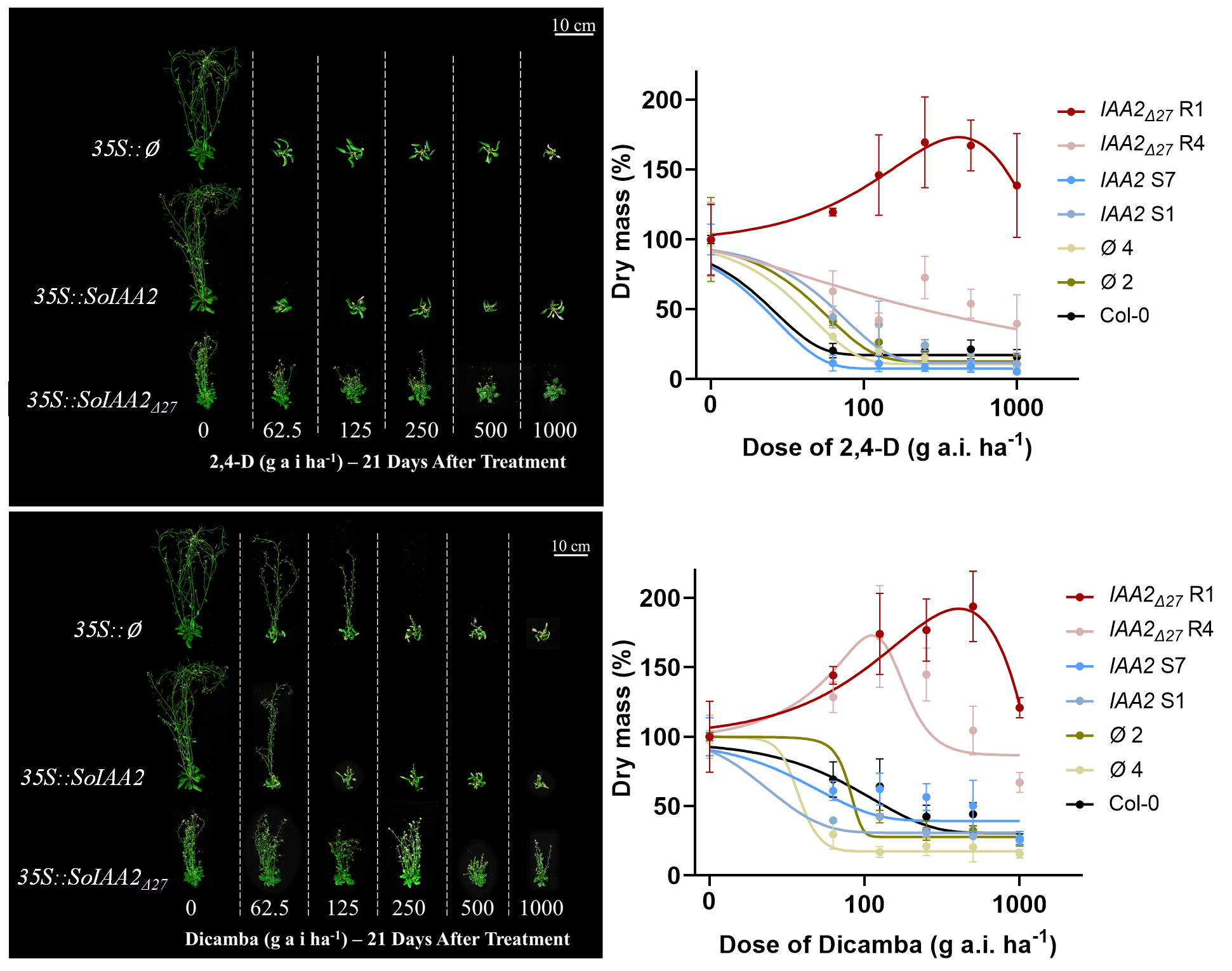

**Supplementary Figure 4. 2,4-D and dicamba dose responses in Col-0 and transformed lines containing null vector, *SoIAA2*, and *SoIAA2_Δ27_*.** Pre-flowering plants (14 d after germination) were treated with doses of 0, 62.5, 125, 250, 500, and 1000 g ai ha^-1^ of 2,4-D and dicamba. Plants were harvested 21 d after herbicide application, dry mass was measured, and dose response models were generated considering untreated plants as 100% of dry mass accumulated (n=3, error bars = SEM).

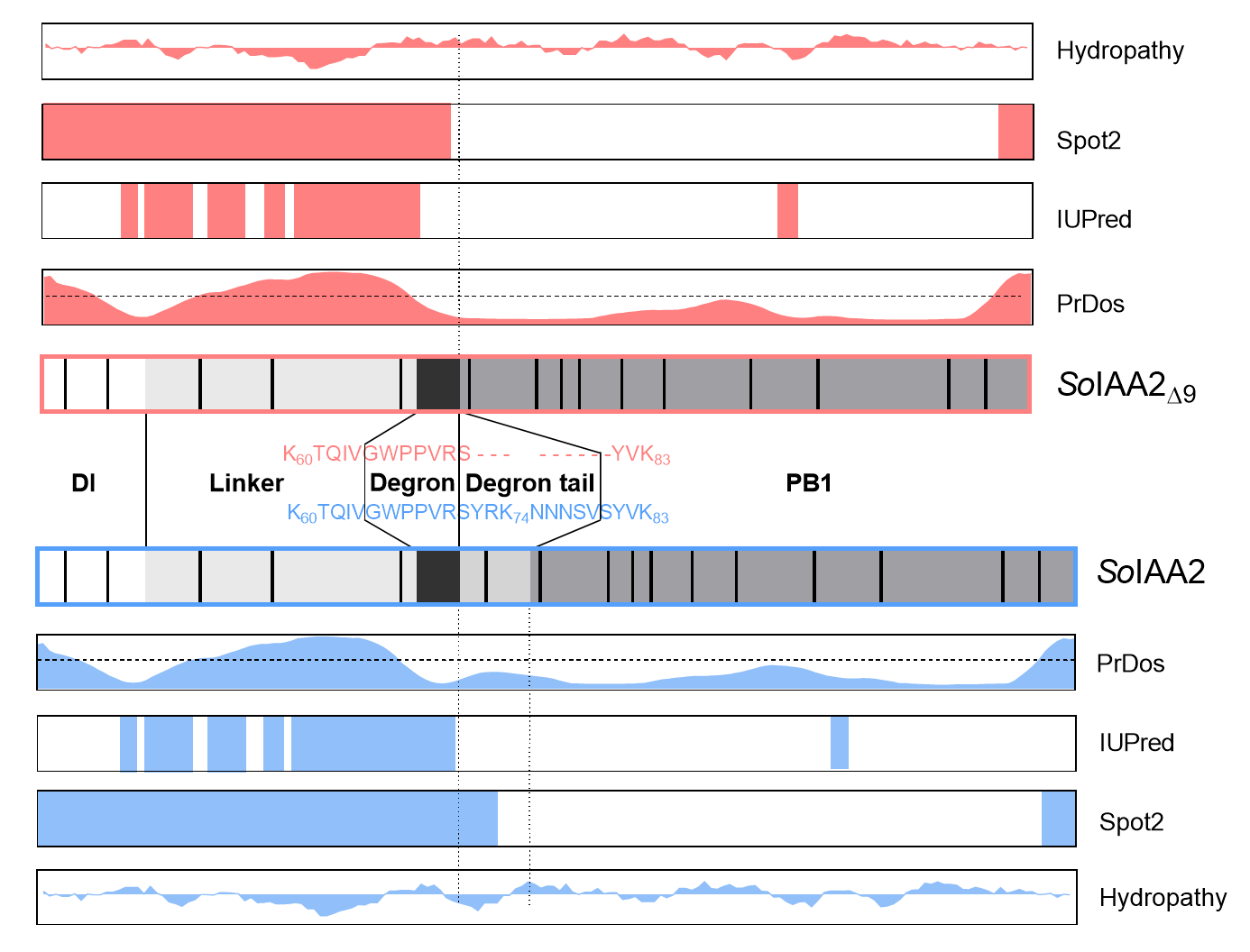

**Supplementary Figure 5. Intrinsic disordered regions (IDR) and hydropathy predictions of *So*IAA2 (blue) and *So*IAA2_Δ9_ (red).** *In silico* analysis of each protein using PrDos, IUPred, and Spot2. PrDos shows disorder predictions (disordered: > 0.6; intermediate disordered: 0.4-0.6; ordered: < 0.4), and the dotted line corresponds to a value of 0.5. IUPred and Spot2 maps show colored regions corresponding to high scores for IDR. Kyte-Doolittle hydropathy maps scaled from -4 to +4, with negative values corresponding to hydrophilic sequences and positive to hydrophobic sequences.

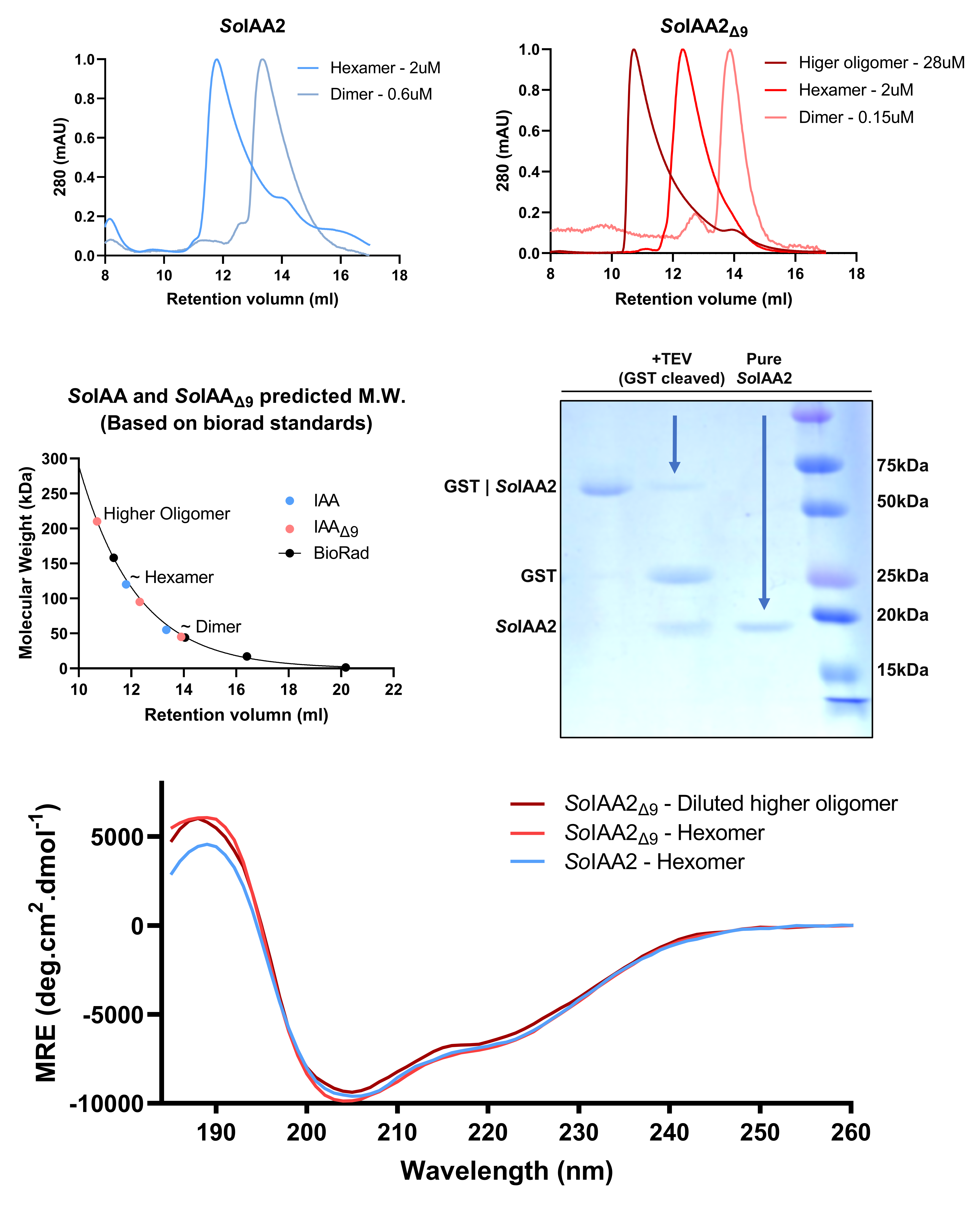

**Supplementary Figure 6. Size exclusion chromatography and CD of recombinant untagged *So*IAA2 and *So*IAA2_Δ9_.** A) SEC Chromatograms for repeated injections from peak elutions for *So*IAA2 (blue) and *So*IAA2_Δ9_ (red) normalized to 1 mAU for clarity. Each reinjection of the peak dilutes the protein ~10 fold and shifts the elution peak, reflecting a change in oligomeric state proportional to the protein concentration. The dark lines correspond to *So*IAA2 and *So*IAA2_Δ9_ at predicted states of approximately 6 and >10 molecules, respectively. Each subsequent dilution and injection results in an eventual shift to a dimeric state for both proteins. B) Plot showing retention volumes for Bio-Rad standards and each of the SEC injections. The standards are fit to a single exponential and used to predict the molecular weight for each protein at various oligomeric states. C) SDS-PAGE showing an example of the proteins at different stages of purification. The first two lanes are the GST fused *So*IAA2 protein (~47 kDa) before and after TEV cleavage. Lane 3 is the final pure protein and lane 4 is the molecular weight standard. GST is noted at ~27 kDa and final pure *So*IAA2 ~20 kDa. D) CD data for *So*IAA2 and *So*IAA2_Δ9_ from the primary peak (dark SEC lines) for each protein, as well as an off-peak sample for *So*IAA2_Δ9_ at 12.7 mL corresponding to an approximate tetramer. All three CD spectra show an alpha, beta, other content of 12.9%: 38.4%: 48.7%, which agrees with a partial structure from the PB1 domain and about half of the protein being disordered or “other”.
